# Contribution of climate models and APSIM phenological parameters to uncertainties in spring wheat simulations: application of SUFI-2 algorithm in northeast Australia

**DOI:** 10.1101/2021.01.28.428676

**Authors:** Brian Collins, Ullah Najeeb, Qunying Luo, Daniel K.Y. Tan

## Abstract

We used SUFI-2 for the first time to calibrate the phenology module of the APSIM-wheat model for 10 spring wheat cultivars cultivated in northeast Australia (south-eastern Queensland). Calibration resulted in an average RMSE of 5.5 d for developmental stages from stem elongation up to flowering. Projections from 33 climate models under the representative concentration pathway 8.5 were used for simulations at 17 sites. Using adapted sowing times, we simulated significantly shorter crop cycles and grain yield improvements for the period 2036-2065 relative to 1990-2019 for three selected cultivars (Hartog, Scout and Gregory). Photoperiod and vernalisation sensitivities were shown to be the largest and smallest contributors to total uncertainties in the simulated flowering day and grain yield. Uncertainties in climate models had a relatively minor contribution to the total uncertainties in the simulated values of target traits. This contribution significantly increased when climate change impact on the target traits was estimated.

## 1 Introduction

Crop phenology, i.e. knowledge of crop developmental phases, is essential for precision farming and crop management. It is a key component of crop models that use cultivar-specific parameters to simulate growth, development and yield of a crop. There has been an increased application of process-based crop models to address the interactive impact of genotype × environment × management interactions (G×E×M) on crop yield and development under a changing climate (e.g. Zheng et al., 2012; Lobell et al., 2015; Stöckle et al., 2018; Webber et al., 2018; Hunt et al., 2019; Liu et al., 2019, 2020; Ababaei and Najeeb, 2020). Crop models have also been extensively adopted in climate change impact and adaptation studies across Australia (Yang et al., 2014; Luo et al., 2018; Hunt et al., 2019; Ababaei and Najeeb, 2020; Liu et al., 2020) and worldwide (Ababaei and Ramezani Etedali, 2017; Stöckle et al., 2018; Liu et al., 2019; Asseng et al., 2015; Challinor et al., 2009a; Pörtner et al., 2014; Rosenzweig et al., 2014). However, due to the complex nature of biophysical factors associated with crop production processes, many uncertainties originate while simulating the impact of the future climate on crops. Sources of uncertainty include projection of greenhouse gas emissions, projection of global warming, projection of local climate change, estimation of crop model parameters and crop model structure (Zhang et al., 2019; White et al., 2011; Challinor et al., 2013; Wallach and Thorburn, 2017).

Previous studies quantified the effect of these uncertainties on estimates of climate change impact on crop production. For example, Luo et al. (2005) and Tao et al. (2008) used the Monte Carlo technique to quantify and manage uncertainties from climate change projections. Studies by Iizumi et al. (2009) and Zhang et al. (2019) used the Markov Chain Monte Carlo (MCMC) technique for examining the probability distribution of biophysical parameters. Asseng et al. (2013), Araya et al. (2015) and Wang et al. (2017) reported a greater contribution of crop model structure to the total uncertainty than general circulation models (GCMs). In contrast, greater uncertainties in crop yield projections from GCMs than those from crop model structure were reported by Tao et al. (2009), Kassie et al. (2015) and Zhang et al. (2019). These conflicting findings show the necessity of assessing each source of uncertainty with the target model(s), within the target study area, and for the target crop(s).

Among these sources, crop model parameter estimation (the so-called crop model calibration or inverse modelling) can be an important source of uncertainty. Calibration is a critical step in developing and applying a simulation model, which uses observational data to estimate unknown model parameters for a better fit of model outputs (Seidel et al., 2018). Common goodness-of-fit criteria for determining crop parameters include visual assessment and statistical/optimisation approaches such as maximum likelihood, ordinary least square, D-statistics (Willmott, 1982), and Bayesian analysis. Some calibration algorithms also generate useful statistical information such as parameter sensitivities for supporting calibration. Compared to direct measurement of parameters in the field, which is time and cost consuming, inverse modelling generates inferences from common measured phenological and production components. Recently, there has been an increased interest in using inverse modelling for calibrating crop models (e.g., Ababaei et al., 2014; Andarzian et al., 2015; Yuan et al., 2017; Hussain et al., 2018; Zhang et al., 2019; Gao et al., 2020). However, most of the previous studies explored the impact of uncertainties in model parameters on the target outputs by assigning arbitrary ranges of variations to a number of parameters (e.g. Bert et al., 2007; DeJonge et al., 2012; Zhao et al., 2014), usually based on literature or previous experiences. This approach could lead to unrealistically large impacts on selected outputs as it does not account for the uncertainty in observations and the interaction between the parameter uncertainty and model structure uncertainty under local conditions.

Soil and crop models have a large array of biophysical and physiological parameters. For such complex models, a sensitivity analysis helps to identify the importance of each parameter to the response of target output variables (Richter et al., 2010). Sensitivity analysis has so far been applied to different cropping systems and climate scenarios to evaluate the importance of input parameters for target outputs using agro-hydrological models (Asseng et al., 2004; DeJonge et al., 2012; Kumar et al., 2014; Zhao et al., 2014). Asseng et al. (2004) showed complex interactions between phenological and physiological traits in high and low yielding environments for wheat yield. The model outputs could be sensitive to both individual parameters and their combinations (Pogson et al., 2012) and the sensitivities of parameters depend on model complexity, the number of crop parameters included in the analysis and the environment (Richter et al., 2010).

Hence, the objectives of present study were to: (1) calibrate phenology module of the widely used Agricultural Production Systems sIMulator (APSIM)-wheat model with a modified version of the SUFI-2 (Sequential Uncertainty Fitting, ver. 2; Abbaspour et al., 2004) calibration and uncertainty analysis algorithm, (2) examine sensitivities of the selected phenological parameters under current and future climate scenarios, (3) assess the impact of parameter uncertainty on the simulated phenology and grain yield and also on the quantification of the impact of climate change on these traits, and (4) evaluate the impact of climate change on wheat crops in northeast Australia.

## 2 Materials and methods

### 2.1 Field experiments

Eight field experiments were conducted over 2 years (2018–2019) at three different locations across southern Queensland, Australia (Table 1). Randomised block design trials with two times of sowing (TOS) and four replicates were conducted each year. Zadoks growth stage (Zadoks et al., 1974) was recorded from stem elongation (Z31) up to flowering (Z65) for all genotypes. Phenology data were recorded as an average of the whole plot (i.e. at least 50% of culms in the plot). Wheat crops were planted late in the cropping season, to ensure they would receive high temperatures during different developmental phases. For example, the crop was planted on July 3 and 9 (conventional sowing) and August 31 and September 3 (late sowing) at The University of Queensland Research Farm, Gatton (27°34’50”S 152°19’28”E). The experiments were also conducted at the Hermitage Research Station, Warwick (28°12’40”S, 152°06’06”E) and Tosari Crop Research Centre, Tummaville (27°54’60”S, 151°30’0”E) with sowing on July 16 (conventional sowing), and September 12 and September 6 (late sowing) in 2018 and 2019, respectively. All the experiments were planted using a cone seeder with a target population of 100 plants m^−2^ and a row-spacing of 250 mm within 5 m × 1 m plots. The crops were irrigated at sowing and cultivated under non-limiting fertiliser conditions. Standard crop management practices including weed, disease and pest control were adopted.

**Table 1.** Field experiments used for calibrating the phenological parameters.

| Location | Coordinates |  | Previous crop | Sowing dates | Year | Irrigation |
| --- | --- | --- | --- | --- | --- | --- |
|  | S | E |  |  |  |  |
| University of Queensland Research Farm, Gatton | 27° 34' 50" | 152° 19' 28" | Fallow | 03-Jul 31-Aug | 2018 | Full |
|  |  |  |  | 09-Jul 03-Sep | 2019 | Full |
| Hermitage Research Station, Warwick | 28° 12' 40" | 152° 06' 06" | Fallow | 16-Jul 12-Sep | 2018 | Supplementary |
| Tosari Crop Research Centre, Tummaville | 27° 54' 60" | 151° 30' 0" | Cotton | 16-Jul 06-Sep | 2019 | Supplementary |

Ten commercial Australian bread wheat (*Triticum aestivum* L.) cultivars with contrasting phenology and adaptation were used in this study (Table 2). These include three high-performing spring cultivars with a wider adaptation to Australian environments i.e. Suntop (mid-season maturity), Mace (early to mid-season maturity) and Scout (mid-season maturity).

**Table 2.** List of the cultivars with varying maturity type and adaptation.

| Cultivar | Maturity type | Adaptation | Reference |
| --- | --- | --- | --- |
| Suntop | Mid maturing | Widely adapted to Australian environments | Graham et al., 2015 |
| Mace | Mid to slow maturing | Widely adapted to Australian environments | Graham et al., 2015 |
| Scout | Mid maturing | Widely adapted to Australian environments | Graham et al., 2015 |
| EGA Gregory | Mid to slow maturing | Nematodes resistance | Graham et al., 2015 |
| EGA Wylie | Mid to slow maturing | Fusarium crown rot resistance | Zheng et al., 2014 |
| Seri-82 | Mid to slow maturing | Drought tolerance | Christopher et al., 2008 |
| Drysdale | Mid to slow maturing | Drought tolerance | Condon et al., 2012; Tausz-Posch et al., 2012 |
| Spitfire | Fast to mid maturing | High grain protein | Graham et al., 2015; Brill et al., 2013 |
| Hartog | Fast to mid maturing | Drought susceptible | Brennan et al., 1991; Christopher et al., 2008 |
| Janz | Fast to mid maturing | Straw strength and standability | Brennan et al., 1991 |

### 2.2 Calibration and uncertainty analysis algorithm

In this study, SUFI-2 calibration and uncertainty analysis algorithm was adopted for calibration of APSIM-wheat crop model (version 7.10). To the best of our knowledge, the present study is the first to use this algorithm for calibrating a crop model. The original SUFI-2 has been implemented in SWAT Calibration and Uncertainty Programs (SWAT-CUP; Abbaspour et al., 2007a; Abbaspour, 2015), which is designed for calibration of Soil and Water Assessment Tool (SWAT). SUFI-2 accounts for the uncertainties in observations while estimating parameter values and performs uncertainty analysis at the same time as parameter estimation. A modified version of SUFI-2 (hereafter, ‘SUFI-2M’) was implemented in a customised package in the R programming environment (R Core Team, 2017).

A step-by-step description of SUFI-2 has been presented by Abbaspour et al. (2007b). All steps and related equations are presented in the Supplementary Material. Briefly, initial uncertainty ranges are allocated to the selected parameters for the first round of sampling. These ranges are subjective and are selected based on literature or previous experience. Then, a Latin Hypercube (LH; McKay et al., 1979) sampling is carried out leading to *LHn* (here, 200) parameter combinations, which should be relatively large. The model is then run *LHn* times and the target traits are stored. A goal function (here, root mean square error, RMSE) is calculated and the sensitivity matrix is created. Next, an equivalent of a Hessian matrix is calculated (Abbaspour et al., 2007b). Then, an estimate of the lower bound of the parameter covariance matrix (*C*) is made (Press et al., 1992) using variance of the objective function values resulting from the *LHn* runs. The estimated standard deviation and 95% confidence interval of each parameter is calculated from the diagonal elements of the covariance matrix. As all parameters are allowed to change, the correlation between any two parameters is quite small and can be evaluated with the diagonal and off-diagonal terms of *C*.

SUFI-2 calculates the 95% prediction uncertainties (95PPU) for all the variables in the objective function (i.e. values of target traits). It is calculated by the 2.5^th^ and 97.5^th^ quantiles of the cumulative distribution of simulated points. The aim is to encapsulate as many measured data as possible within the 95PPU band (P-factor: percentage of observed data that fall within the 95PPU), and to reduce average distance between the upper and the lower 95PPU (d-Factor: the degree of uncertainty). The ‘ideal’ outcome is that 100% of the measurements fall within the 95PPU range and d-Factor is close to zero (Abbaspour et al., 2007b). This ideal situation is generally not achievable. We seek to see most of observations fall within the 95PPU range. At the same time, we prefer to have a small 95PPU range (i.e. uncertainty range). No specific recommendation exists for these two factors, like any other goodness of fit measure. However, a value of >70% can be suggested for P-factor while having R-factor of around 1 is acceptable (Abbaspour et al., 2007b).

Parameter ranges are updated at the end of each iteration. This approach ensures that the updated parameter ranges are always cantered around the best estimates. It is recommended that of the highly correlated parameters, those with smaller sensitivities should be fixed to their best estimates and removed from additional sampling rounds (Abbaspour et al., 2007b). For the present study, number of iterations was set to 100.

A few modifications were introduced to the original SUFI-2 routine. First, updating parameter ranges in SUFI-2M is performed at the beginning of each iteration instead of at the end. This way, we assure that the finally selected ‘best’ parameter set has been chosen from a range of which the corresponding 95PPU range brackets a pre-defined percentage of observations (here, 70%). If the updating is performed at the end of each iteration, final parameter ranges may or may not satisfy this criterion. In the modified version, parameter ranges are not updated if no improvement in the goal function has been achieved. Moreover, only the best parameter set is used to update the parameter ranges, instead of using the average of the top *p* (a user-defined number) best parameter sets, which is the case in the original version.

Another modification was the introduction of a ‘boosting’ option. With this option, *LHn* is reduced at the beginning of each iteration (see Supplementary Material for the equations). This option reduces *LHn* proportionally to changes in parameter ranges and makes the optimisation procedure significantly faster.

### 2.3 Calibration setup

Four phenological parameters were selected for calibration based on previous experiences on model performance: (1) vernalisation sensitivity (vern_sens), (2) photoperiod sensitivity (photop_sens), (3) thermal time from ‘end of juvenile’ to ‘floral initiation’ (tt_end_of_juvenile), and (4) thermal time from ‘floral initiation’ to ‘flowering’ (tt_floral_initiation). To evaluate the capability of SUFI-2M for crop model calibration, the selected phenological parameters were calibrated for 10 selected spring wheat cultivars. Following a common and widely accepted approach (e.g. Chenu et al., 2011; Hammer et al., 2014; Lobell et al., 2015; Zheng et al., 2016; An-Vo et al., 2018; Ababaei et al., 2019), most crop parameters were assumed to be similar across cultivars and therefore equal to the default values for the base cultivar (i.e. Hartog). Observed phenology data from Gatton 2018 (second sowing) and Gatton 2019 (first sowing) experiments were used for validation (i.e. ~35% of the observations) each representative of a number of trials in terms of heat-shock and drought patterns. The rest of the data were used for calibration. In order to consider the uncertainty in observations and account for its impact on parameter estimation, all replications were used for calibration as independent measurements.

### 2.4 Simulation setup

The APSIM-wheat model (Keating et al., 2003; Holzworth et al., 2014), which has been widely tested and used in Australia (e.g. Asseng et al., 2001; Lilley and Kirkegaard, 2007; Hochman et al., 2009; Chenu et al., 2011; Christopher et al., 2016; Wang et al., 2018; Ababaei and Chenu, 2019, 2020; Ababaei and Najeeb, 2020), was adopted (version 7.10) to simulate wheat crop growth and development under current and future climate scenarios. Heat-shock and frost impacts were estimated using methods described by Lobell et al. (2015), Ababaei and Chenu (2020) and Zheng et al. (2015a).

Out of ten calibrated spring wheat cultivars, three cultivars (Hartog, Scout and Gregory) of contrasting maturity habits and different ranges of parameter uncertainties were chosen for simulations with sowing dates between April 1 and July 31 at 7-day intervals. At each location, soil characteristics, initial soil nitrogen, fertilisation levels and planting density were set to represent local soils and farming practices. (see Table 1 in Chenu et al., 2013). Soil initial conditions were reset on January 1 each year to the median level based on long-term simulations (Chenu et al., 2013). A small amount of irrigation was applied at sowing, if needed, to raise soil moisture of the top layer to the lower limit of plant-extractable soil water so that seeds could germinate the day after sowing.

### 2.5 Climate data

Historical daily weather data, including maximum and minimum temperature, solar radiation and rainfall, were obtained from the SILO patched point dataset (http://apsrunet.apsim.info/cgi-bin/silo; Jeffrey et al., 2001) for the period 1976-2019 at 17 selected sites across northeast Australia (Figure 1; Ababaei and Chenu, 2020). Locations were selected to represent the wheat-producing regions of Queensland and New South Wales (Chenu et al., 2013). Monthly projections of precipitation and minimum and maximum temperatures from 33 GCMs for future period centred on 2050 were obtained from the Coupled Model Intercomparison Project 5 (CIMIP5; Taylor et al., 2012). Future climate scenarios were constructed for 2036-2065 (hereafter, the ‘2050’ climate) by downscaling to a daily time step. Downscaling was performed by applying projected changes in local monthly means to the daily temperature and rainfall data for the period of 1976-2005 (Lobell et al., 2015). The 30-year period of 1990-2019 (hereafter, the ‘2005’ climate) was simulated as the benchmark scenario to quantify the impact of climate change.

**Figure 1.**
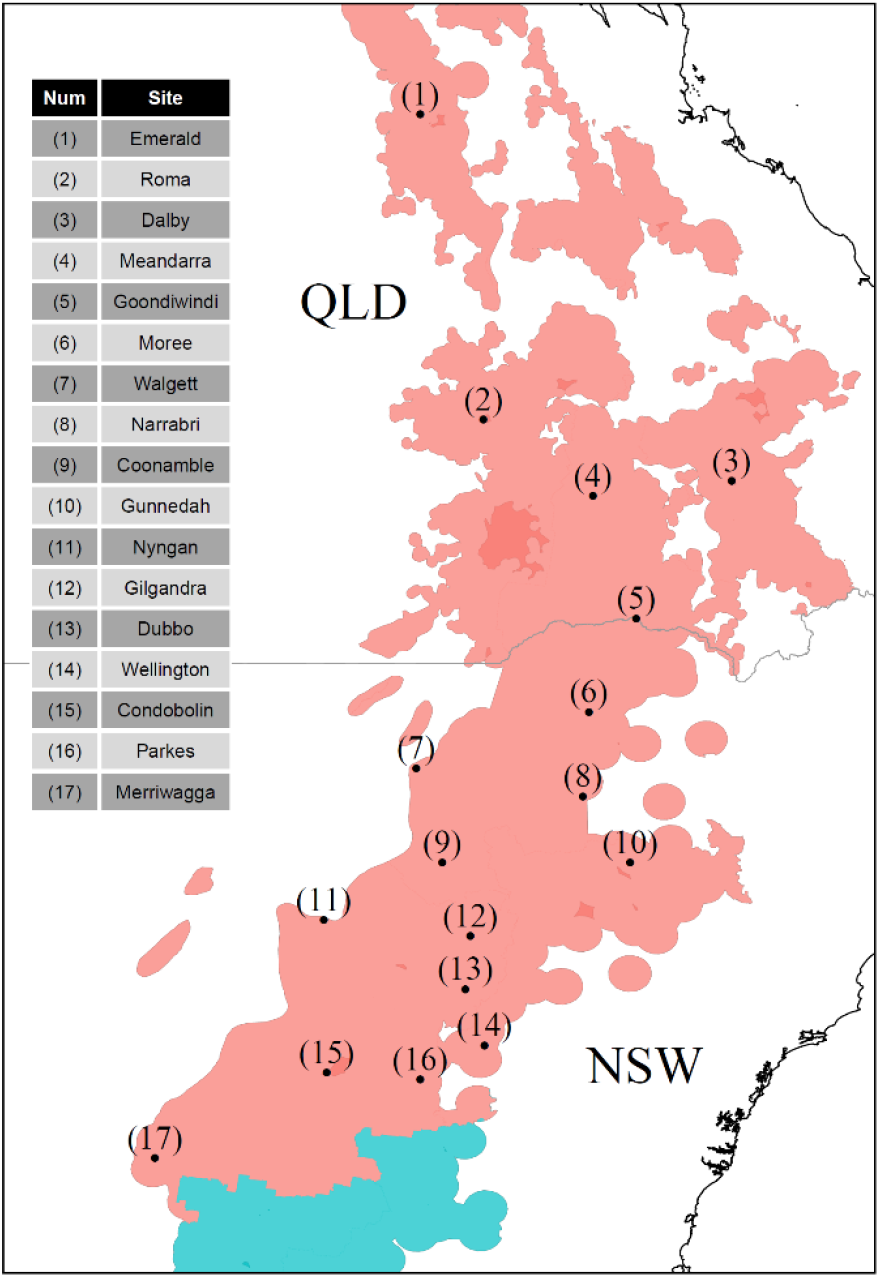
The 17 selected sites representative of the major wheat-producing regions in northeast Australia (depicted with red shading).

Atmospheric [CO_2_] was set at 541 ppm for the 2050 climate, as projected by the representative concentration pathway (RCP) 8.5, which assumes ‘business as usual’ CO2 emission. For the 2005 climate, atmospheric [CO_2_] data was obtained from Ziehn et al. (2016). In APSIM-wheat, elevated [CO_2_] linearly increases transpiration efficiency from a cultivar-specific reference value at 350 ppm by 37% when [CO2] reaches 700 ppm (Reyenga et al., 1999). It is also related to the radiation use efficiency (RUE) which is adjusted with a temperature response function. In APSIM-wheat, RUE at 20oC increases by 14% when [CO_2_] increases to 541 ppm (Lobell et al., 2015).

### 2.6 Optimum sowing dates

The ‘best’ sowing date in each season, i.e. leading to the highest grain yield, was used to quantify the target traits (flowering date and grain yield) under each climate scenario. This was done to minimise the effect of suboptimal management choices and give a more realistic evaluation of the ‘net’ impact of climate change on wheat crops when cultivated on the best sowing date in each season. In order to investigate the impact of climate change on optimum sowing dates, sowing windows were determined separately under each climate scenario. For the 2050 climate, one common sowing window was identified for all the GCMs. The windows were determined as the period between 25^th^ and 75^th^ quantiles of the selected dates over the 30 simulated seasons.

### 2.7 Quantifying the impact of parameter uncertainty on simulated traits

Any uncertainty in estimated values of crop parameters may lead to substantial deviations in the simulated values of target traits from the values simulated by the ‘best’ calibrated parameter sets. Therefore, after calibration was performed and the final parameter ranges were determined for each cultivar, a set of parameter values (hereafter, ‘uncertain’ parameter sets) was created for each cultivar to analyse the impact of parameter uncertainties on the simulated values of the target traits (flowering day and grain yield). To that end, the lower and upper bounds and the mid-point of each parameter range were chosen. Each combination of these three values and the four selected parameters (3^4^ = 81 parameter sets) was considered as an individual ‘virtual’ cultivar. For each parameter, the ‘uncertain’ parameter sets represent the range of uncertainty around the ‘best’ parameter value and allow quantification of the impact of uncertainties on the simulated traits.

We evaluated the contributions of model parameters and climate models to the total uncertainty of target traits using an Analysis of Variance (ANOVA; Tao et al., 2018; Zhang et al., 2019). Variance components were estimated as the corresponding contribution (in percentage) of each factor to the total sum of squares. ‘Cultivar uncertainty’ was estimated as the variation of target traits across the ‘virtual’ cultivars while the simulated target traits were averaged across the GCMs. ‘GCM uncertainty’ was estimated as the variation of target traits across the 33 GCMs while averaged across the ‘virtual’ cultivars.

### 2.8 Sensitivity analysis of APSIM-wheat phenological parameters

APSIM-wheat sensitivities to uncertainties in wheat phenological parameters were evaluated with a global sensitivity analysis (GSA) approach (Abbaspour et al., 2007b; Faramarzi et al., 2010; Abbaspour, 2015). GSA overcomes the drawbacks of local optima (or one-at-a-time) approaches by exploring the entire multi-dimensional parameter space simultaneously, leading to a better quantification of the influence of each parameter and the interactions between parameters (Saltelli et al., 2008). A multiple linear regression system, which relates the parameters generated by the Latin Hypercube sampling to the objective function values (i.e. the values of target traits) in all simulated seasons, was separately constructed for each season as well as for all simulated seasons together. A t-test was used to identify the significance of each estimated parameter value. The sensitivity indices (SI) calculated by this method are estimates of the average changes in the target traits (here, flowering day and grain yield) stemming from uncertainties in each parameter while all other parameters are also changing. That is, the SIs are based on linear approximations and only provide partial information about the sensitivity of the objective function to model parameters.

Sensitivity analysis was independently performed under the 2005 and 2050 climates. To quantify the contribution of climate models to the total uncertainty in parameter sensitivities, sensitivity analysis was performed for each GCM separately as well as for all GCMs collectively under the 2050 climate. As the range of selected parameters were different, each parameter was scaled (by subtracting the average and dividing by the standard deviation) before fitting the multiple linear regressions. To obtain a better understanding of the magnitude of sensitivities, we further compared the contributions of location, climate models and inter-annual variability to the total variance of target traits using ANOVA.

## 3 Results

### 3.1 Parameter calibration

Calibration was performed with the aim of encapsulating at least 70% of the observed phenology data within the uncertainty range of model simulations (95PPU). Figure 2 shows the observed vs simulated phenology for the 10 wheat cultivars calibrated with SUFI-2M along with the simulation uncertainty ranges. The overall root mean square error (RMSE) of the calibration/validation phase was 5.6/5.5 d (Figure 2). Calibration significantly improved model performance compared with when the APSIM default parameters were adopted (Supplementary Material Figure S1). Considering all the individual replications, the lowest and highest errors were related to *cv* Hartog (RMSE = 3.9 d; Figure 3a), and *cv* Seri-82 and Drysdale (RMSE = 7.1 d), respectively. Figure 3b presents the ‘best’ calibrated value for phenological parameters along with the uncertainty ranges of the parameters.

**Figure 2.**
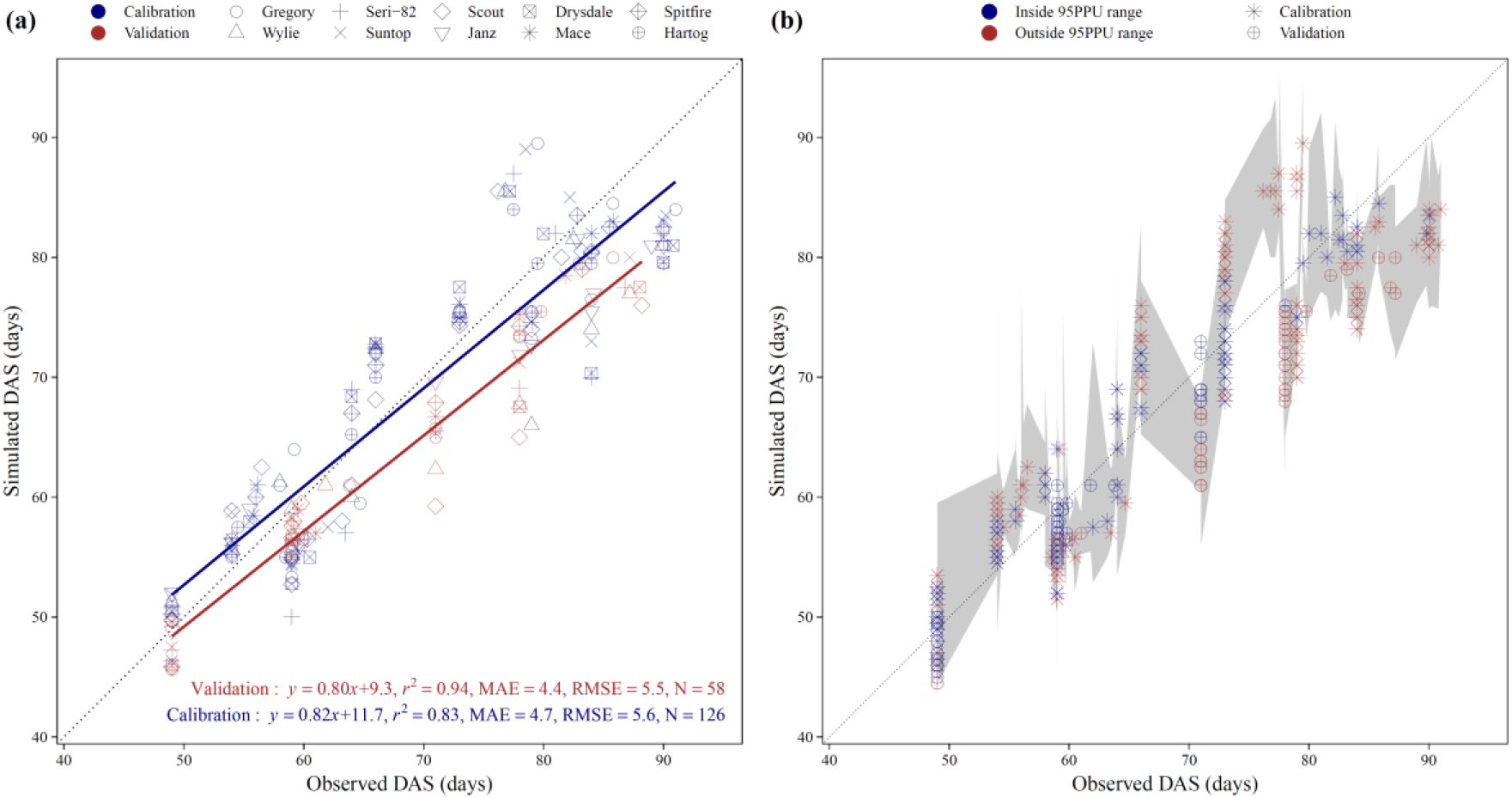
APSIM-wheat calibration results with the SUFI-2M algorithm across three locations in northeast Australia (south-eastern Queensland): **(a)** calibration for the 10 selected spring wheat cultivars; **(b)** uncertainty ranges in the simulated phenology along with the observed phenology for all the individidual replications. Data were averaged across replications in panel **(a)**. RMSE is the root mean square error, MAE is the mean absolute error and N is the number of data points. Observations included Zadoks growth stages from stem elongation (Z31) up to flowering (Z65). ‘Inside’ and ‘outside’ refer to the 95% prediction uncertainties (95PPU) range and whether the observed values fell within this range or not.

**Figure 3.**
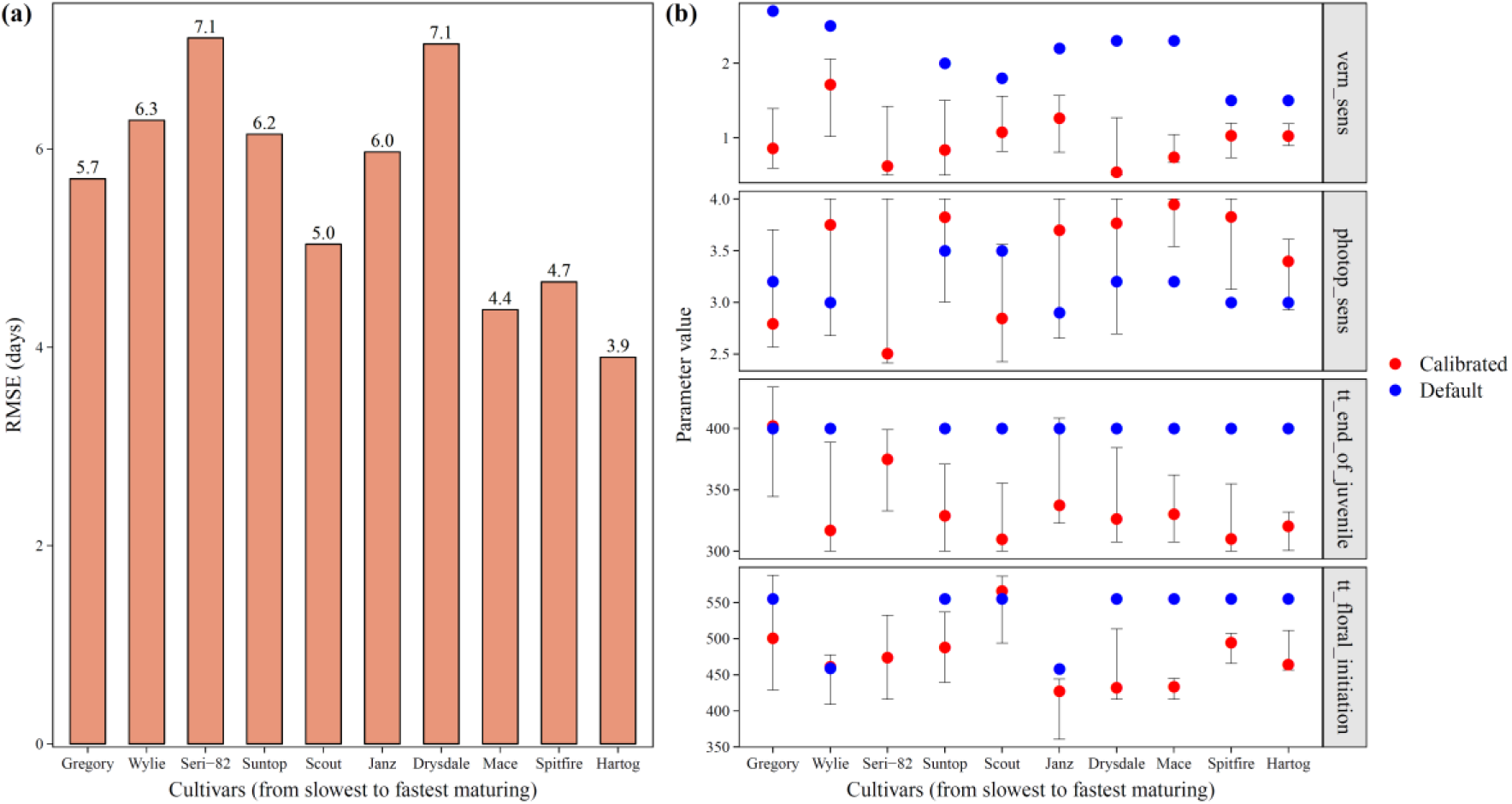
Root mean square error (RMSE) for the 10 selected spring wheat cultivars after calibration, considering all the individual replications **(a)**, and the ‘best’ calibrated values (red points) and uncertainty ranges (error bars) of each phenological parameter along with the default values of APSIM cultivar-specific parameters (blue points) **(b)**. Parameters included: (1) vernalisation sensitivity (vern_sens), (2) photoperiod sensitivity (photop_sens), (3) thermal time from ‘end of juvenile’ to ‘floral initiation’ (tt_end_of_juvenile), and (4) thermal time from ‘floral initiation’ to ‘flowering’ (tt_floral_initiation).

### 3.2 Climate change is expected to shift sowing dates to earlier dates in the season

Sowing windows were determined under each climate scenario separately for three spring wheat cultivars (Hartog, Scout and Gregory) of contrasting maturity habits (Figure S2). Under the 2050 climate and all the studied sites, except Emerald (for all cultivars) and Nyngan (only for *cv* Hartog), the start and mid-point of the sowing windows are expected to occur earlier in the season than in 2005. The shift was estimated to be 6.7, 7.0 and 7.9 d for the start date and 9.0, 9.8 and 11.9 d for the mid-point for Hartog, Scout and Gregory, respectively. Moreover, the sowing windows would be 7.5, 6.4 and 6.7 d shorter than those in 2005.

### 3.3 Crop model sensitivity changes over time and in space

Sensitivity of APSIM-wheat to uncertainties in crop phenological parameters varied over time (Figure 4; Supplementary Material Figures S3-S4) and in space (Supplementary Material Figure S5). For example for *cv* Hartog, approximately >92% of the total variance in the sensitivities of simulated flowering day and grain yield to changes in vernalisation sensitivity (vern_sens) was explained by residuals (i.e. inter-annual variations), which implies a high temporal variability in the SIs (Figure 5). These variance components were estimated to be larger in 2050 than in 2005, and for flowering day than for grain yield. Other parameters were more variable across the sites though the variance components were generally smaller in 2050 than in 2005 and larger for grain yield than for flowering day. Contribution of climate models to the total variance of the SIs was markedly smaller (<1%) than other sources of variance. On the other hand, 9-33% of the total variance in the sensitivity of flowering day to uncertainties in phenological parameters in 2050 was explained by location, while this component was generally smaller for grain yield.

**Figure 4.**
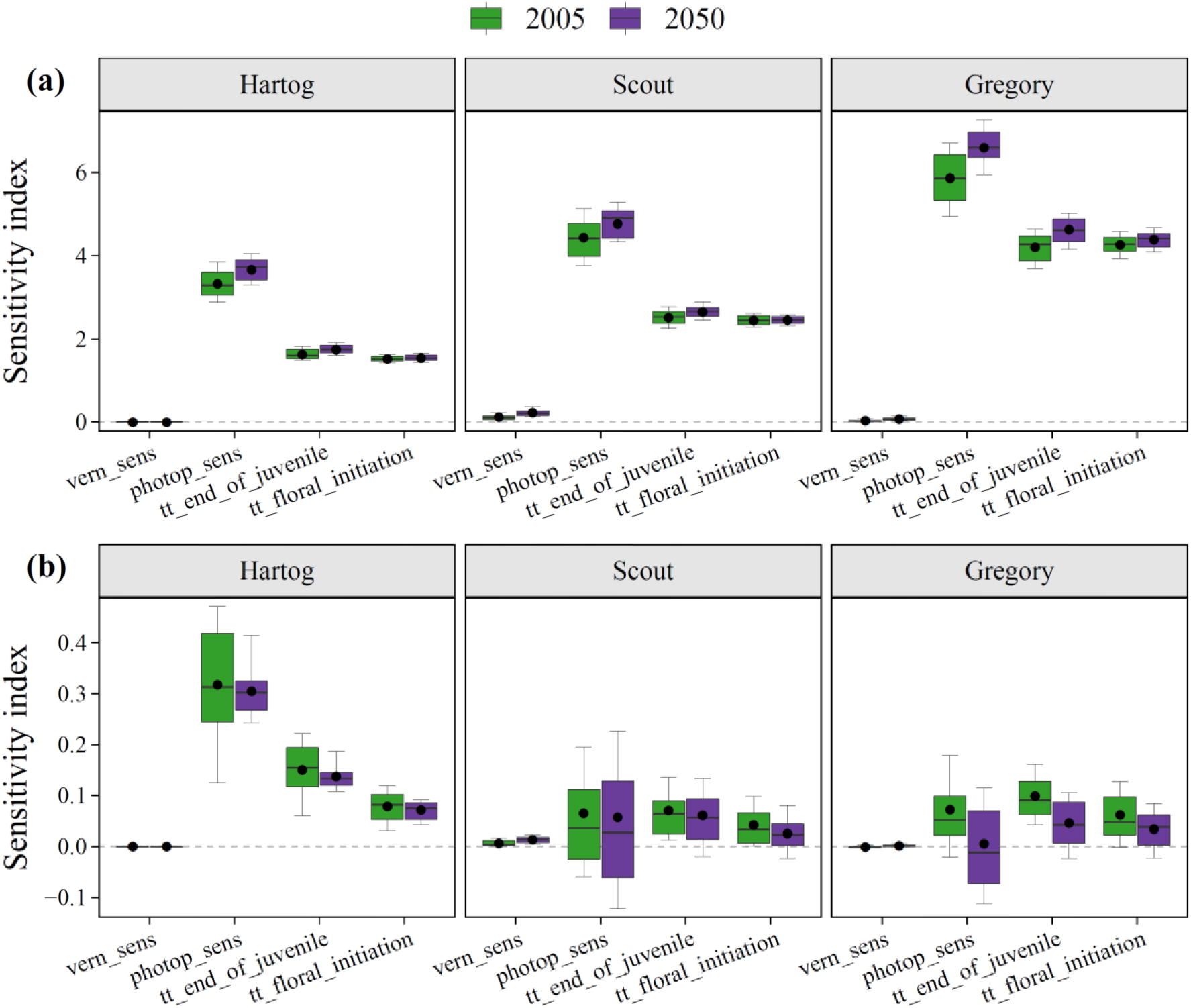
Sensitivity indices (SIs, averaged across 30 seasons) of the selected four phenological parameters for flowering day **(a)** and grain yield **(b)** under the 2005 (current) and 2050 (future) climates for three selected spring wheat cultivars across 17 sites in easutern Australia. Box plots show 10, 25, 50, 75 and 90th quantiles along with the means (black points). Parameters included: (1) vernalisation sensitivity (vern_sens), (2) photoperiod sensitivity (photop_sens), (3) thermal time from ‘end of juvenile’ to ‘floral initiation’ (tt_end_of_juvenile), and (4) thermal time from ‘floral initiation’ to ‘flowering’ (tt_floral_initiation).

**Figure 5.**
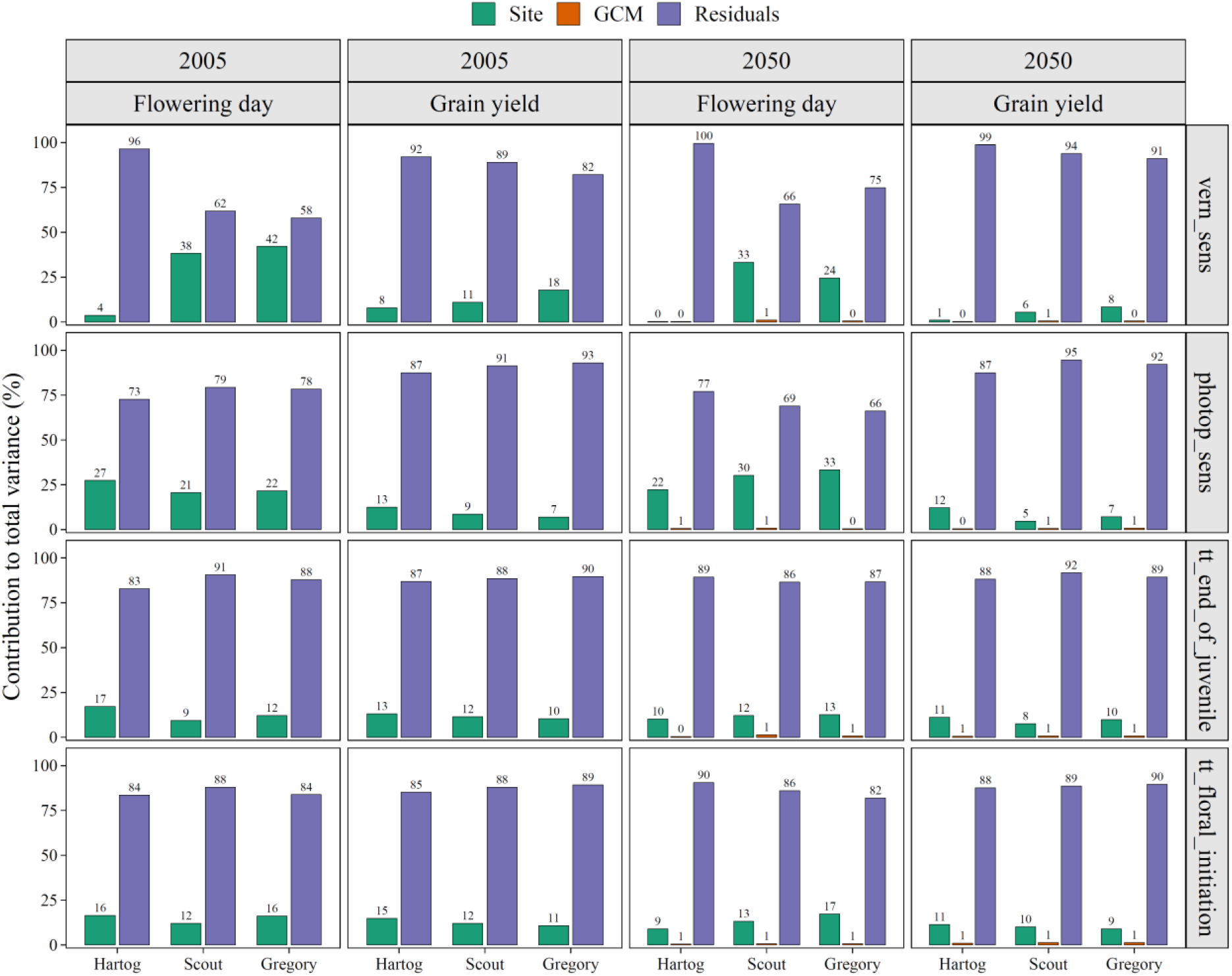
Variance components of the sensitivity of the target traits (flowering day and grain yield) to the four phenological parameters under the 2005 (current) and 2050 (future) climates for three selected spring wheat cultivars. Parameters included: (1) vernalisation sensitivity (vern_sens), (2) photoperiod sensitivity (photop_sens), (3) thermal time from ‘end of juvenile’ to ‘floral initiation’ (tt_end_of_juvenile), and (4) thermal time from ‘floral initiation’ to ‘flowering’ (tt_floral_initiation).

### 3.4 Crop parameter sensitivities are expected to change in the future

For the four phenological parameters, we calculated the SIs related to wheat flowering day and grain yield under both climate scenarios (Figure 4). Among the calibrated parameters, uncertainties in photoperiod sensitivity (‘photop_sens’) was the most influential source of uncertainty on the simulated flowering day (Figure 4a), while vernalisation sensitivity (‘vern_sens’) was the least influential. Gregory, a slow-maturing spring cultivar, was most sensitive to uncertainties in ‘photop_sens’, ‘tt_end_of_juvenile’ and ‘tt_floral_initiation’ among the studied cultivars. These sensitivities are expected to slightly increase under future climate scenario, especially for ‘photop_sens’.

The order of most sensitive parameters for grain yield was the same as for phenology (Figure 4b) for Hartog, however, the SIs were less variable across the selected sites for Scout and Gregory, except for ‘vern_sens’. Unlike for flowering day, it is expected that the sensitivity of phenological parameters decreases in 2050 relative to 2005, though the shifts were generally insignificant for Hartog and Scout.

The only non-significant (*P*>0.05; i.e. *P* value of the t-test) SIs were related to ‘vern_sens’ for flowering day and across a few of the sites (Roma, Dalby, Meandarra and Walgett) for grain yield (Supplementary Material Figure S5). Sensitivity of flowering day to uncertainties in all the four parameters, except for ‘vern_sens’, generally increased from north to south (in the southern hemisphere) and the trend was strongest for ‘photop_sens’ under both climate scenarios. A similar but relatively weaker trend was observed for grain yield, though the spatial patterns were more heterogeneous.

### 3.5 Parameter uncertainties affect trait quantifications

Larger uncertainties in phenological parameters of *cv* Scout and Gregory, as compared with Hartog, led to larger deviations of simulated flowering day with uncertain parameter sets from the values simulated with the ‘best’ parameter sets (Figure 6a), suggesting larger sensitivities. The mean absolute errors (MAE) were estimated to be <1 d for Hartog under both climates and 3.9 and 4 d for Gregory under the 2005 and 2050 climates, respectively. However, these deviations were not directly reflected in the simulated grain yields (Figure 6b). While the uncertainty ranges for phenological parameters were larger for Scout than for Hartog, the simulations with uncertain parameter sets (representative of parameter uncertainties) resulted in smaller deviations from the simulations with the ‘best’ parameter set (a normalized MAE of 11% for Scout vs 15% for Hartog in 2005). This suggests that the simulated grain yield is more sensitive to other environmental and management factors than flowering day is.

**Figure 6.**
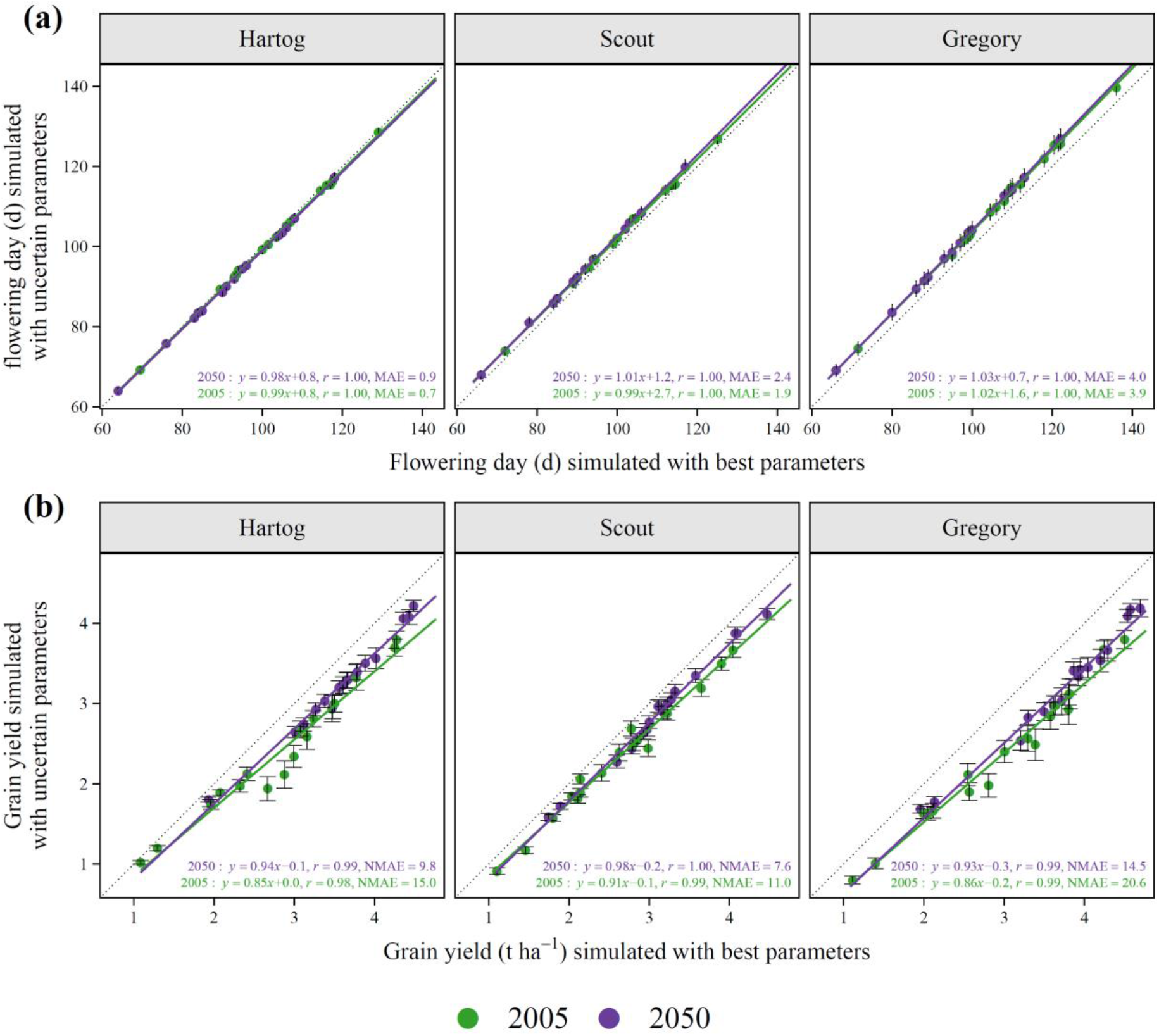
Flowering day **(a)** and grain yield **(b)**, averaged across 30 seasons, simulated with the best (x-axis) and uncertain (y-axis) parameter sets (averaged across 81 parameter combinations) for three selected spring wheat cultivars under the 2005 (current) and 2050 (future) climate scenarios. A larger deviation of y-axis values from x-axis values suggests a larger sensitivity.

### 3.6 Climate change is expected to enhance phenology and improve grain yield in northeast Australia

Wheat crops under the warmer and drier climate of 2050 were projected to reach flowering significantly earlier as compared with 2005 (Figure 7a). Using the best (**|** uncertain) parameter sets for three selected cultivars and at a regional scale, it was shown that Hartog, Scout and Gregory would reach flowering day 9.9 (**|** 10.2), 9.5 (**|** 9.1) and 11.6 (**|** 11.1) d earlier in 2050 than in 2005. While this impact may reduce the time available for biomass assimilation, the wheat grain yield could still benefit from the elevated atmospheric [CO_2_] levels, leading to grain yield increases by 22.8 (**|** 30.1), 20.4 (**|** 24.3) and 25.1 (**|** 32.5)% for the studied cultivars planted on the optimum sowing dates (Figure 7b, Supplementary Material Figure S6). The results showed that the inclusion of parameter uncertainties led to statistically significant deviations from the trait values simulated with the ‘best’ parameter sets (*P*<0.05) for all the tested cultivars.

**Figure 7.**
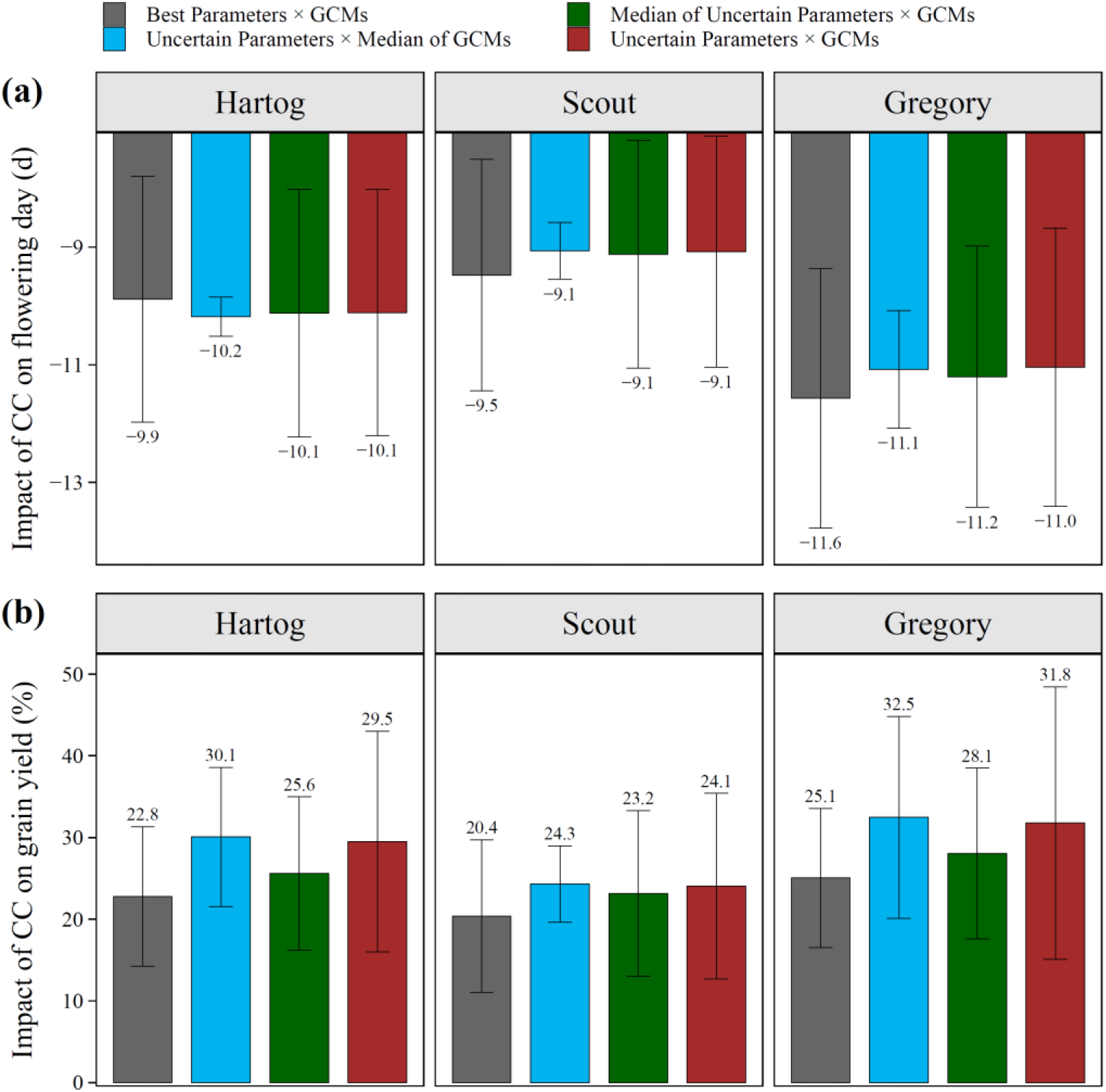
The impact of climate change (CC) on flowering day **(a)** and grain yield **(b)** at regional scale (i.e. averaged acorss 30 seasons and 17 sites) simulated with the best and uncertain parameter sets for three selected speing wheat cultivars. Error bars show standard deviation of the simulations across 33 GCMs (gray, green and red bars) and 81 uncertain parameter sets (blue and red bars).

With the best sowing time adopted in each season under each climate scenario, it is anticipated that all the studied cultivars would experience a significant improvement in grain yield level in 2050 relative to 2005 (Supplementary Material Figure S6), except in Emerald. Across all other locations, it would be expected that Hartog, Scout and Gregory have higher grain yields by 9.7-63.1%, 5.4-53% and 9.9-67.5%, respectively. Emerald was the only site that is expected to experience a reduction in average grain yield in 2050, at which Hartog, Scout and Gregory were expected to have lower simulated grain yields by 1.4, 2.8 and 2.5%, respectively.

Considering all the site × GCM combinations (Figure 8), 99.6, 99.4 and 99.9% of combinations showed a negative shift in flowering day in 2050 for Hartog, Scout and Gregory, respectively. For grain yield, in 95.5, 93.3 and 94.6% of the combinations a positive impact of climate change was simulated for the three cultivars.

**Figure 8.**
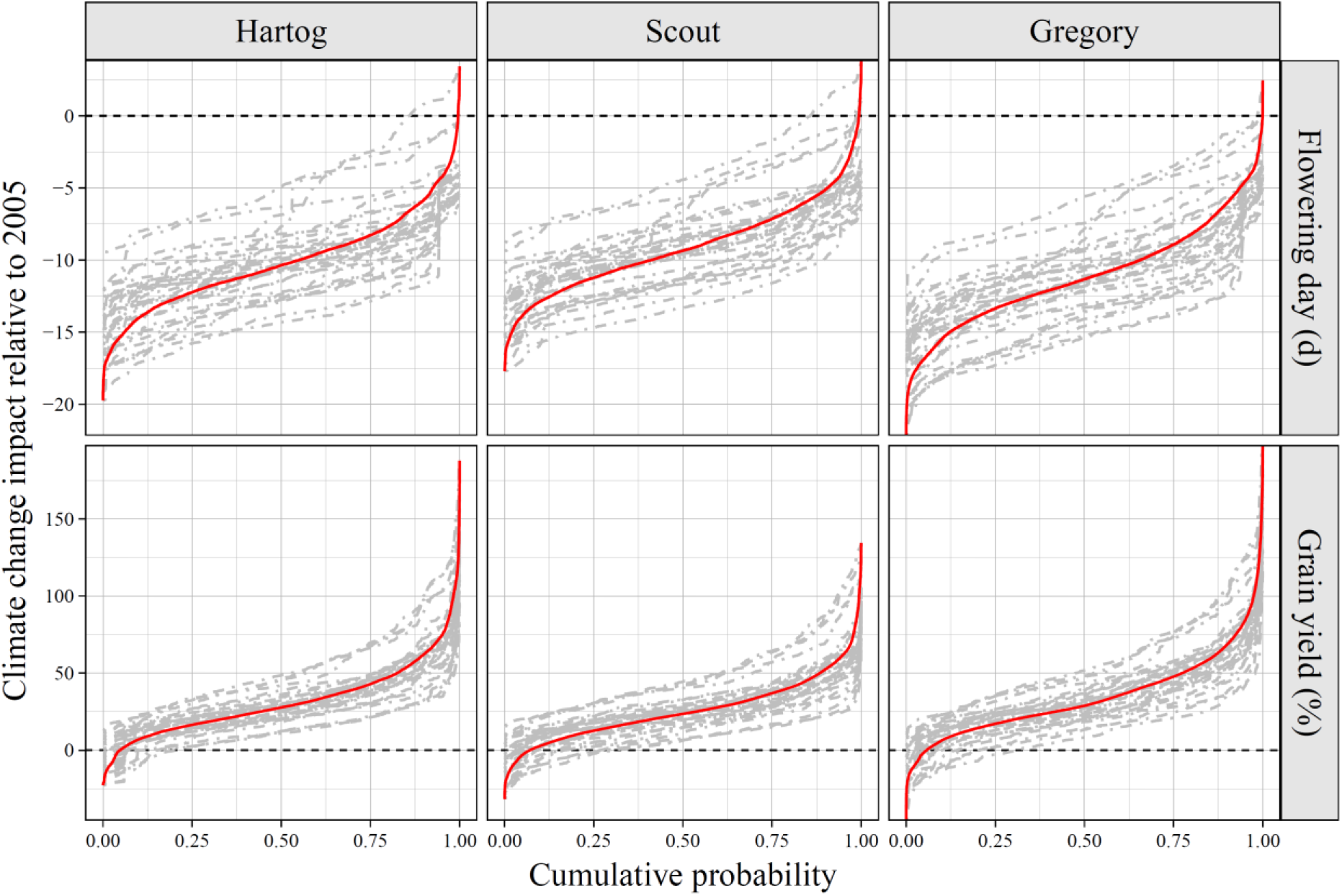
The cumulative probability of the simulated impact of climate change on flowering day (absolute change, d) and grain yield (relative change, %) with each of the 33 GMCs (gray lines) and all the GCMs combined (red line) for three selected speing wheat cultivars. Cumulative probabilities on the horizontal axis show the probability of observing impacts less than the numbers on the vertical axis.

### 3.7 Contribution of crop parameters and climate models to simulation uncertainties

Both crop parameters and climate models (GCMs) contribute to the uncertainties in the target traits. Regarding absolute values of the target traits under the 2050 climate, the contribution of crop parameters to uncertainties was larger than climate models (upper panels in Figure 9; Figure 10) while residuals (i.e. inter-annual variability) were the largest variance component. The contribution of crop parameters to the total uncertainties in the simulated flowering day was larger for Gregory and Scout than for Hartog, while this component was the smallest for grain yield of Scout. The contribution of phenological parameters (|climate models) to the total uncertainties were estimated to be 26 (|9), 38 (|7) and 55 (|5)% for the simulated flowering day and 15 (|8),10 (|8) and 16 (|7)% for the simulated grain yield for Hartog, Scout and Gregory, respectively.

**Figure 9.**
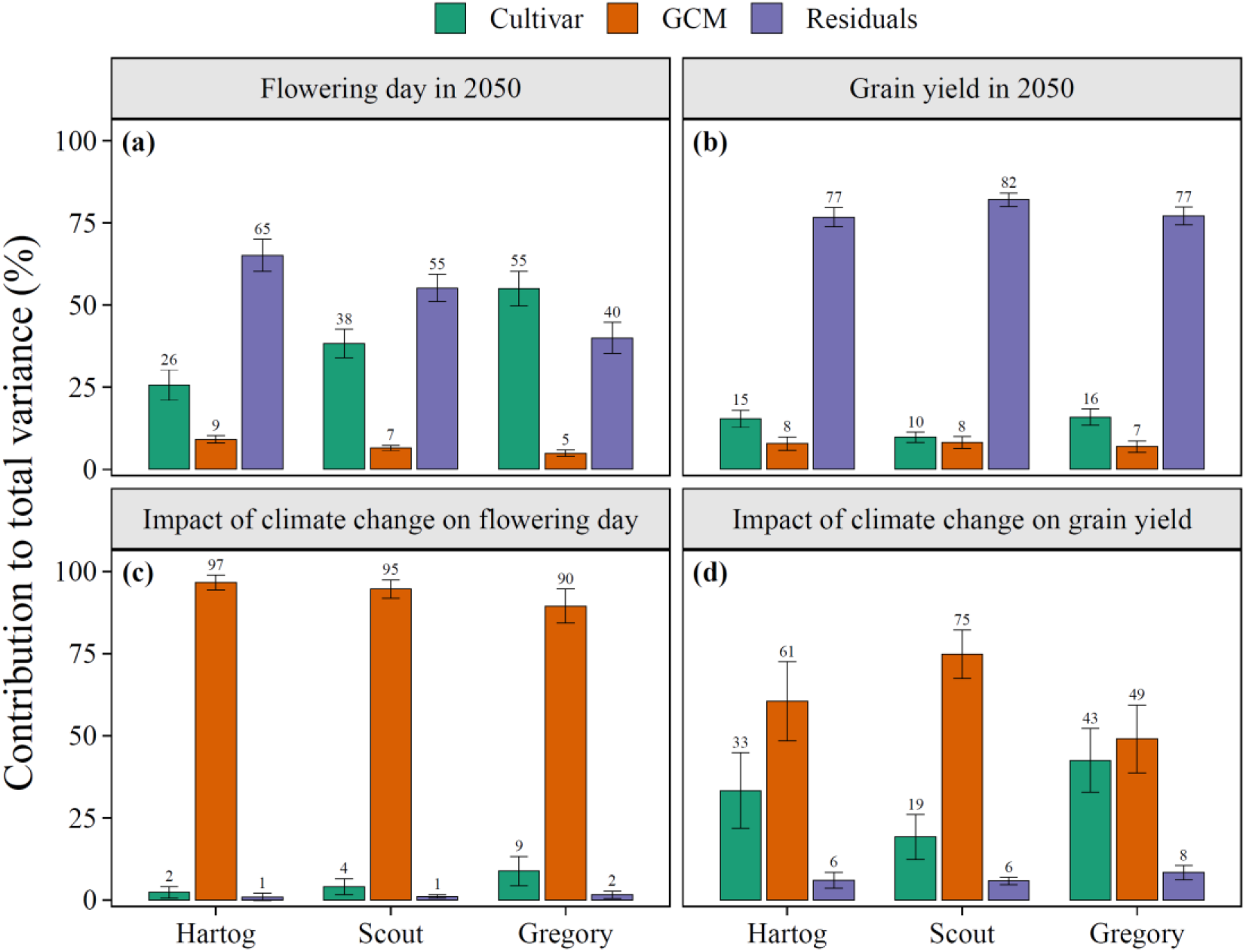
Variance components of the simulated values of the target traits (flowering day and grain yield) under the 2050 climate **(a-b)** and of the simulated impact of climate change on each trait **(c-d)** for the three selected cultivars. Values were averaged across the 17 selected sites in northeast Australia.

**Figure 10.**
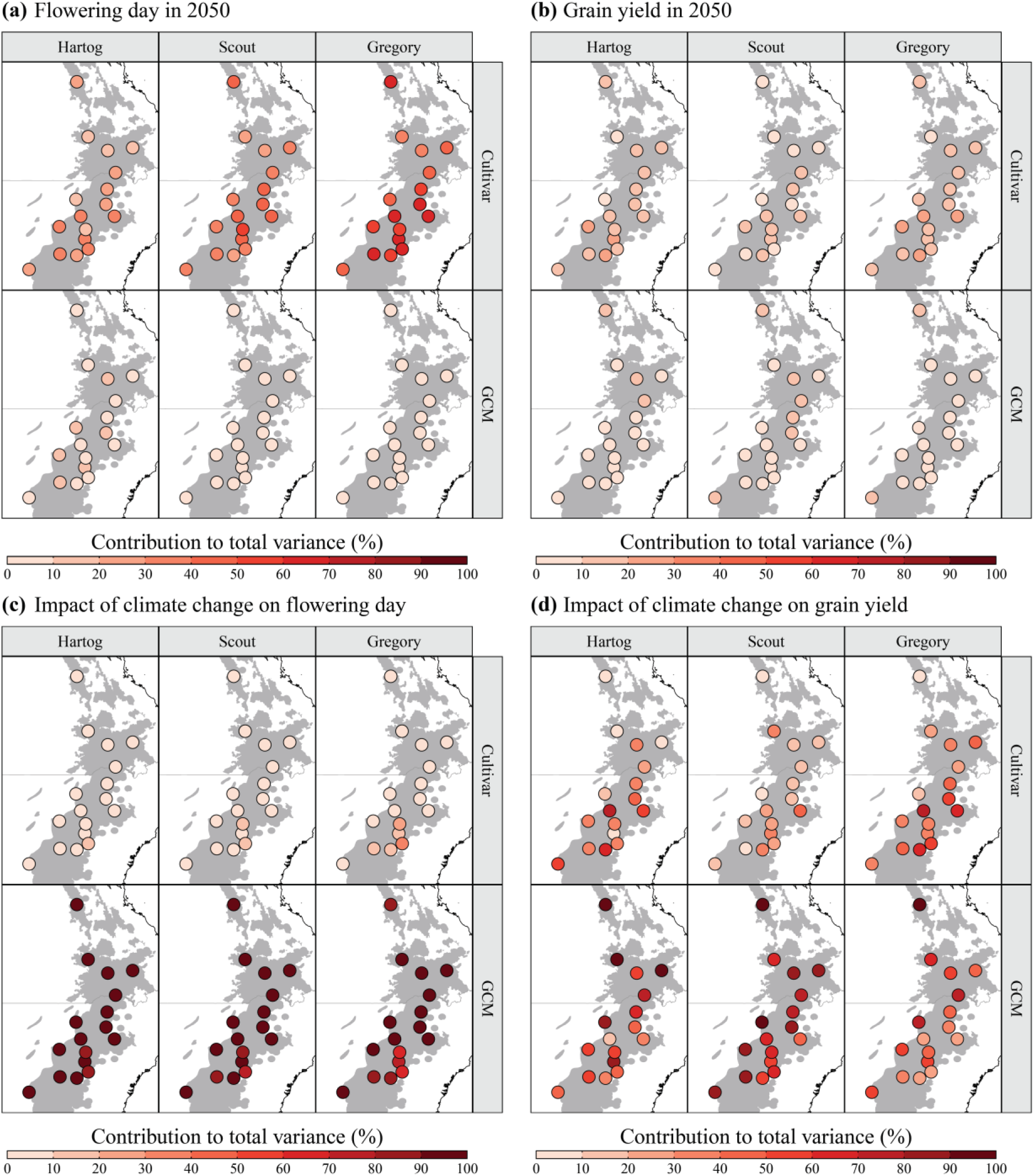
Variance components of the simulated values of the target traits (flowering day and grain yield) under the 2050 climate **(a-b)** and of the simulated impact of climate change on each trait **(c-d)** for the three selected spring wheat cultivars across 17 selected sites in northeast Australia.

On the other hand, the contribution of climate models to the total uncertainties in the simulated impact of climate change (i.e. change in target traits in 2050 relative to 2005) on flowering day and grain yield was significantly larger than crop parameters, especially for flowering day (lower panels in Figure 9; Figure 10). Gregory (with the widest parameter uncertainty ranges) and Hartog (with the narrowest parameter uncertainty ranges) had the lowest and highest contributions of phenological parameter to the total variances, respectively.

### 3.8 Spatial pattern of sensitivity indices in northeast Australia

Contributions of the two major sources of uncertainty to the total uncertainty in simulated flowering day and grain yield and the impact of climate change on target traits were significantly correlated with latitude (Figure 10). Considering all three cultivars, the correlation coefficient was estimated to be −0.21 and 0.24 for flowering day, −0.41 and 0.51 for grain yield, −0.24 and 0.27 for the impact of climate change on flowering day and −0.45 and −0.51 for the impact on grain yield, respectively.

Figure 11 shows the impact of climate change on flowering day and grain yield simulated using the ‘best’ parameter sets against the averages of the simulations with the uncertain parameter sets. Considering all the uncertainties in the selected parameters, the impact of climate change on flowering day was simulated with a MAE of 0.2, 0.5 and 0.6 d for Hartog, Scout and Gregory, respectively. These deviations were smaller than the deviations estimated for the absolute values of the target traits under each climate scenario (Figure 6). The same was the case for grain yield, with MAE values estimated to be 7, 4.3 and 7.6%. This implies that APSIM-wheat is more sensitive to uncertainties in phenological parameters than uncertainties in climate models when the aim is to quantify the absolute values of target traits under a future climate scenario. On the contrary, the model is more sensitive to uncertainties in climate models when the target is to ‘compare’ the target traits under current and future climate scenarios.

**Figure 11.**
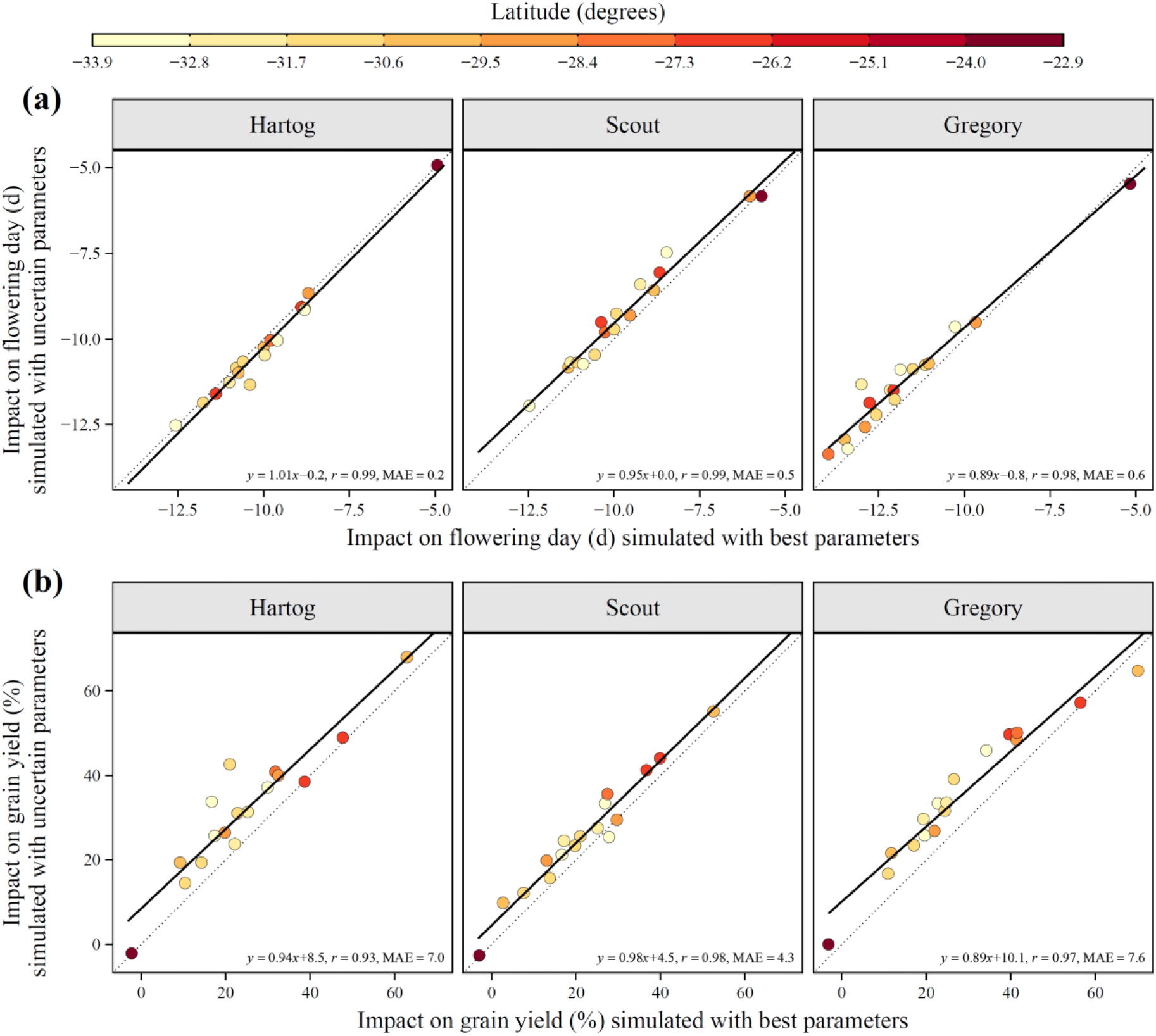
The impact of climate change on flowering day **(a)** and grain yield **(b)** of three selected spring wheat cultivars. Values are averaged across 30 seasons, simulated with the best (x-axis) and uncertain (y-axis) parameter sets (averaged across 81 parameter combinations). A larger deviation suggests a larger sensitivity.

## 4 Discussion

### 4.1 Calibration

We successfully calibrated the APSIM-wheat crop model in northeast Australia (south-eastern Queensland) using SUFI-2M algorithm. SUFI-2M resulted in a robust calibration of the model with an overall RMSE of 5.5 d (3.9-7.1d) despite a considerable variations in the observed phenology of the 10 selected wheat cultivars. While more detailed data on wheat growth stages were used in this study than just data on heading, flowering or maturity days, these deviations are considerably lower or at least comparable to the reported deviations by other similar studies, for example, the reported RMSE for *cv* Janz was 6.2 d for days to heading (Zheng et al. 2012), 6.2 d for flowering day (Flohr et al. 2017), 7-8 d for flowering and maturity days of spring wheat cultivars (Liu et al. 2018), 1.4–7.2 days for days to heading in barley (Liu et al. 2020) and 9.4-35.3 d for days to flowering in rice (Gao et al. 2020).

In previous studies, other parameter estimation routines have been used for calibrating hydrological and crop models. For example, Gao et al. (2020) applied three commonly used calibration methods to the CSM-CERES-Rice phenology model of the Decision Support System for Agrotechnology Transfer (DSSAT), including Ordinary Least Square (OLS), MCMC, and Generalized Likelihood Uncertainty Estimation (GLUE). They reported that selection of the calibration routine had implications for parameter estimates and uncertainty quantifications and found MCMC more reliable than GLUE in quantifying model uncertainty. Iizumi et al. (2009) and Tao et al. (2009) applied the MCMC technique to crop models for paddy rice and spring maize to optimize a set of regional-specific parameters and quantified the uncertainty of yield estimation associated with model parameters. They found that MCMC is a powerful technique to optimize multiple parameters, to quantify their uncertainties and to investigate the impacts of climate variability on crop productivity. As each of these techniques have been used with different crop or hydrological models, in different locations and for different purposes/crops, it is not practically possible to make a thorough comparison. Therefore, a choice should be made based on availability of statistical knowledge, computation power and the objectives of the study.

### 4.2 Sensitivity analysis

Our findings showed that flowering day was mainly sensitive to uncertainties (i.e. changes) in photoperiod sensitivity (photop_sens) and thermal time from ‘end of juvenile’ to ‘floral initiation’ (tt_end_of_juvenile) for all the three studied spring cultivars (Figure 4). The sensitivity of these parameters did not only depend on the maturity habit of a cultivar, but on the location (i.e. latitude) and time (i.e. sowing year; Supplementary Material Figures S3-S4). For grain yield, the ranking of parameter sensitivities was the same as for flowering day for *cv* Hartog. However, uncertainties in ‘tt_end_of_juvenile’ and thermal time from ‘floral initiation’ to ‘flowering’ (tt_floral_initiation) were the most influential parameters for Scout and Gregory. For all the three selected cultivars, vernalisation sensitivity (vern_sens) was the least sensitive parameter for the simulated flowering day and grain yield. Our finding stands in contrast to the findings by Zhao et al. (2014) who reported ‘vern_sens’ being the most sensitive parameter among the parameters influential on grain yield at two sites in eastern Australia. This can be explained by the initial ranges of parameter values that were used in that study (0-5 for ‘vern_sens’ and ‘photop_sens’). They chose the ranges based on the range of values for existing cultivars in the APSIM cultivar-specific parameter sets, while the ranges (~0-2 for ‘vern_sens’ and ~2.5-4 for ‘photop_sens’) used in the present study were chosen based on a local calibration with the detailed phenology data and accounting for uncertainties in field observations.

In APSIM-wheat, the length of growing season, especially the vegetative and reproductive phases, determine the amount of biomass accumulation and biomass allocation to grains. Leaf area index (LAI) and dry matter biomass build up quickly until flowering. Vegetative phases are highly sensitive to photoperiod and vernalisation. The thermal time from emergence to end of juvenile is affected by photoperiod and vernalisation sensitivity factors and the number of vernalization days (Zheng et al., 2015b). This implies the importance of sowing time and maturity habit in the quantification of the sensitivity of these parameters.

Grain yield is a more complex trait than flowering day and is sensitive to numerous parameters (e.g. Zhao et al., 2014). Richter et al. (2010) ranked high the phenological development and leaf area dynamics and ranked low the physiological parameters in terms of sensitivity of grain yield to these parameters. They also showed that parameter sensitivities changed in different environments (i.e. sowing time × location). In the present study, we focused on four phenological parameters and the impact of uncertainties in those parameters on grain yield without changing other potentially influential parameters. Therefore, the estimated sensitivity may change in other studies if additional physiological and morphological parameters are included using different environments / cultivars.

### 4.3 Climate change impact on wheat phenology and grain yield

The present study found that wheat flowering will advance by 9 to 12 days by 2050 across the 17 selected sites in northeast Australia (Fig 10a). This is in line with findings by previous studies such as Luo (2016) who simulated 5 to 11 days advancement in the key phenological events of wheat crops by 2050. Similarly, Liu et al. (2018) suggested 7-8 days shortening of sowing-anthesis phase per degree of warming. Wang et al. (2018), Luo et al. (2018), Zheng et al. (2012) and Yang et al. (2014) also reported a substantial boost in wheat phenology under future warmer environments.

It is anticipated that wheat grain yield would potentially increase by 20.4-25.1% in 2050 relative to 2005, depending on the selected cultivar (Fig 11). These numbers must be considered the ‘net’ impact of climate change when the optimum sowing time is chosen in each season. Our results confirm the findings by Ghahramani et al. (2015), Wang et al. (2018) and Hunt et al. (2019) who reported a substantial increase in wheat grain yield under future climates. For example, Ghahramani et al. (2015) simulated an 18% increase in Australia’s national wheat yield by 2030 assuming optimal adaptation. Similarly, Wang et al. (2018) adopted 1961–2000 as the baseline period and reported a 10.7% increase in wheat grain yield across Australia (up to 10% in eastern Australia) by 2050 under RCP 8.5, if autonomous adaptation strategies (i.e. adapted cultivated cultivar and sowing time) would be adopted. Hunt et al. (2019) reported a 0.54 t/ha (~25%) increase in national average wheat yield with early sowing systems combined with slower-developing wheat genotypes.

A negative trend in water-limited yield potential (e.g. 27% between 1990 and 2015 by Hochman et al., 2017) or increased effect of heat shocks have been reported on wheat yield (e.g. 4.6% in each decade by Ababaei and Chenu, 2020). Yang et al. (2014) reported a generally decreasing trend in NSW wheat yield (3.4 to −14.7 %) for the period centred on 2030 when compared to the baseline period of 1961–1990 which is in contrast with our findings. The reason that the estimated yield improvements by 2050 in the current study are higher than some of the previously reported values is that we used 1990-2019 as the baseline scenario instead of 1961-2000 or 1976-2005. That is, our estimates accounts for the already lower yield over the period of 1990-2019 relative to the period of 1961-2000 or 1976-2005. Further, we selected the ‘best’ sowing date in each season, which would lead to the highest grain yield under each climate scenario. Changes in wheat grain yield under future climates depend on location and time of sowing as suggested by Luo (2016) and Luo et al. (2018). The difference in the magnitude of increases in wheat yield between the current study and Anwar et al. (2015) and the change sign opposite to some of previous studies (e.g. Yang et al., 2014) could be attributed to different greenhouse gas emission scenarios, GCMs, locations, time periods and cultivars considered.

### 4.4 Contribution of uncertainty sources

This study investigated the uncertainties arising from the two key sources, i.e., crop model phenological parameters and climate projections. We performed analyses on the simulated values of target traits as well as on the simulated impact of climate change on those traits. We showed a large contribution from crop model phenological parameters to the total uncertainties in the simulated flowering day and grain yield. Holzkämper et al. (2015) stated that the relative importance of uncertainties in climate projections and model parameters depends on local conditions. We can confirm this conclusion as we observed a large spatial variation in the sensitivity indices across the study area.

On the other hand, the uncertainties in climate models are expected to play a more important role than phenological parameters when the aim is to quantify the impact of climate change on target traits (Figure 9). The latter is supported by Tao et al. (2018), who estimated the contribution of crop parameters and GCMs to the total uncertainty in the simulated grain yield to be 42 and 46% at Jokioinen and 24 and 59% at Lleida, Finland (averaged across seven crop models). Studies by Challinor et al. (2009), Kassie et al. (2015) and Zhang et al. (2019) also showed climate projections to be a larger contributor to the total uncertainty in simulations of the impact of climate change on target traits.

The uncertainties in crop model structure was not addressed in the current study, though have been evaluated in previous studies (Tao et al., 2009; Asseng et al., 2013; Araya et al., 2015). Some studies showed that variation in crop model structures could contribute more to the total uncertainty than variation across GCMs while others reported conflicting results. A recent study by Liu et al. (2018) concluded that uncertainty stemming from crop model structure might be larger than crop parameter estimation and the total uncertainty would be larger under a warmer climate due to extra uncertainties from climate projections. These findings and conflicts suggest that the contributions of crop model structure, crop parameter estimation and climate models to the total uncertainty need to be evaluated at specific location(s) and with relevant crop model(s) and crop(s). Moreover, continuing improvement of GCMs and using more robust calculation routines for estimating crop model parameters are necessary to account for uncertainties in field observations and parameter estimation procedure.

## 5 Conclusion

In this study, a slightly modified version of SUFI-2 (SUFI-2M) was used to calibrate the APSIM-wheat model based on two years of experimental data at three locations in northeast Australia (south-eastern Queensland). An overall RMSE of 5.5 d in simulating flowering day of 10 spring wheat cultivars was found in the present study. This suggests that SUFI-2M is a robust and computationally efficient calibration and uncertainty analysis algorithm which can be used for similar research activities.

Sensitivity analyses indicated that uncertainties in photoperiod sensitivity, thermal time from ‘end of juvenile’ to ‘floral initiation’ (tt_end_of_juvenile), and thermal time from ‘floral initiation’ to ‘flowering’ were significantly more influential than uncertainties in vernalisation sensitivity in terms of simulated flowering day and grain yield. We also found a substantial inter-annual variability in parameter sensitivities.

We estimated that climate change would advance wheat phenology by 9.5.-11.6 d at a regional scale across three selected spring cultivars and would have a positive impact on wheat grain yield (20.4-25.1%), if the best sowing date is selected in each season. There are high confidence in the direction of the impact on the target traits, with a probability of shortened time to flowering and positive impact on grain yield estimated to be 99% and 93%, respectively.

We found that variance decomposition of the simulated flowering day and grain yield was significantly correlated with latitude. Uncertainty in the simulated flowering day and grain yield was strongly influenced by the selected crop model parameters than the GCMs, which contributed to the total uncertainty in the simulated values of target traits by less than 9%. On the other hand, the contribution of the uncertainties from the GCMs was the largest component when the impact of climate change on the target traits was to be quantified (>90% for flowering day and 49% for grain yield). It was concluded that the contribution of various sources of uncertainty depended on the environment (i.e. location and sowing time), the maturity habit of the cultivated cultivar, and the target trait.

## 6 Acknowledgement

This research has been funded by the University of Queensland.

