## Supplementary Material for "Contribution of climate models and APSIM phenological parameters to uncertainties in spring wheat simulations: application of SUFI-2 algorithm in northeast Australia"

#### SUFI-2M Algorithm

A step-by-step description of SUFI-2 has been presented by Abbaspour et al. (2007b). Briefly, initial uncertainty ranges (*B**) are allocated to the selected parameters for the first round of sampling. These ranges are subjective and are chosen based on experience. Then, a Latin Hypercube (LH; McKay et al., 1979) sampling is carried out leading to $LH_n$ (here, 200) parameter combinations, which should be relatively large. The model is then run $LH_n$ times and the target traits are stored. Goal function (*G*; here, root mean square error, RMSE) is calculated and the sensitivity matrix (*J*) is created, as follows:

**Eq (1)** $$J_{ij} = \frac{\Delta G_i}{\Delta B_j} \quad i = 1, \ldots, N_2^{LHn} \quad j = 1, \ldots, m$$

Where $B_j$ is parameter *j*, *N* is the number of rows in *J* (i.e. number of all combinations of two simulations), and *m* is the number of parameters. Next, equivalent of a Hessian matrix (*H*) is calculated (Abbaspour et al., 2007b):

**Eq (2)** $$H = J^T J$$

Then, an estimate of the lower bound of the parameter covariance matrix (C) is made (Press et al., 1992) using the variance of the objective function values resulting from the *LHn* runs ($S_G$):

**Eq (3)** $$C = S_G^2 (H)^{-1}$$

The estimated standard deviation and 95% confidence interval of each parameter ($B_j$) is calculated from the diagonal elements of the covariance matrix (c):

**Eq (4)** $$S_j = \sqrt{C_{jj}}$$

**Eq (5)** $$B_{j,lower} = B_j^* - t_{v,0.025} S_j$$

**Eq (6)** $$B_{j,upper} = B_j^* + t_{v,0.025} S_j$$

Here, $B_j^*$ is the parameter *B* from the solution which produces the smallest value of the objective function, and *v* is the degrees of freedom ($LH_n$-*m*). As all parameters are allowed to change, the correlation between any two parameters is quite small. Parameter correlations can be evaluated with the diagonal and off-diagonal terms of *C*, as follows:

**Eq (7)** $$r_{ij} = \frac{C_{ij}}{\sqrt{C_{ii}}\sqrt{C_{jj}}}$$

Assessing the uncertainties in parameters of interest is the next step. SUFI-2 calculates the 95% prediction uncertainties (95PPU) for all the variables in the objective function (*G*). It is calculated by the 2.5$^{th}$ and 97.5$^{th}$ quantiles of the cumulative distribution of simulated points. The aim is to encapsulate as many measured data as possible within the 95PPU band (P-factor: percentage of observed data that fall within by the 95PPU), and to reduce average distance (*d*) between the upper and the lower 95PPU (i.e. the degree of uncertainty):

**Eq (8)** $$\bar{d}_x = \frac{1}{k}\sum_{l=1}^{k}(X_U - X_L)$$

Where *k* is the number of observed data, and $X_U$ and $X_L$ are the upper and lower 95PPU, respectively. The 'ideal' outcome is that 100% of the measurements fall within by the 95PPU range and *d* is close to zero (Abbaspour et al., 2007b). This ideal situation will generally not be achieved. Therefore, a reasonable alternative measure (R-factor) is calculated using the standard deviation of the measured variable ($\sigma_x$), for which a value of <1 is desirable:

**Eq (9)** $$R - factor = \frac{\bar{d}_x}{\sigma_x}$$

Ideally, we would like to see most of observations fall within the 95PPU range. However, we would at the same time like to have a small 95PPU range (i.e. uncertainty range). No hard numbers exist for these two factors, like any other goodness of fit measure. A value of >70% can be recommended for P-factor while having R-factor of around 1 is acceptable. Eventually, parameter ranges are updated (*B'*), as follows:

**Eq (10)** $$B'_{j,min} = B_{j,lower} - \max\left(\frac{B_{j,lower}-B_{j,min}}{2}, \frac{B_{j,max}-B_{j,upper}}{2}\right)$$

**Eq (11)** $$B'_{j,max} = B_{j,upper} + \max\left(\frac{B_{j,lower}-B_{j,min}}{2}, \frac{B_{j,max}-B_{j,upper}}{2}\right)$$

This approach ensures that the updated parameter ranges are always cantered around the best estimates. It is recommended that of the highly correlated parameters, those with smaller sensitivities should be fixed to their best estimates and removed from additional sampling rounds (Abbaspour et al., 2007b). For the present study, number of iterations was set to 100.

A few modifications were introduced to the original SUFI-2 routine that was described by Abbaspour et al (2007b). First, updating parameter ranges in SUFI-2M is performed at the beginning of each iteration instead of at the end. This way, we assure that the finally selected best parameter set has been chosen from a range of which the corresponding 95PPU range brackets a pre-defined percentage of observations (here, 70%). If the updating is performed at the end of each iteration, final parameter ranges may or may not satisfy this criterion. In the modified version, parameter ranges are not updated if no improvement in the goal function has been achieved. Moreover, only the best parameter set is used to update the parameter ranges, instead of using the average of the top *p* (a user-defined number) best parameter sets, which is the case in the original SUFI-2.

Another modification was the introduction of a 'boosting' option. With this option, $LH_n$ is reduced at the beginning of each iteration using the following equations:

**Eq (1)** $$A = \min\left(1, mean\left(\frac{B'_{max} - B'_{min}}{B^*_{max} - B^*_{min}}\right)\right)$$

**Eq (2)** $$LH_{n,min} = \max\,(m + 2, 0.2 \times LH_n^*)$$

**Eq (3)** $$LH_n = \min\,(LH_n, \max\left(LH_{n,min}, ceiling\left(LH_n^* \times \sqrt{A}\right)\right))$$

Where $B^*$ are the initial parameter ranges, $LH_{n,min}$ is the minimum acceptable value for $LH_n$, and $LH_n^*$ is the initial value of $LH_n$. This option reduces $LH_n$ proportionally to changes in parameter ranges and makes the optimisation procedure significantly faster.

### Supplementary Figures

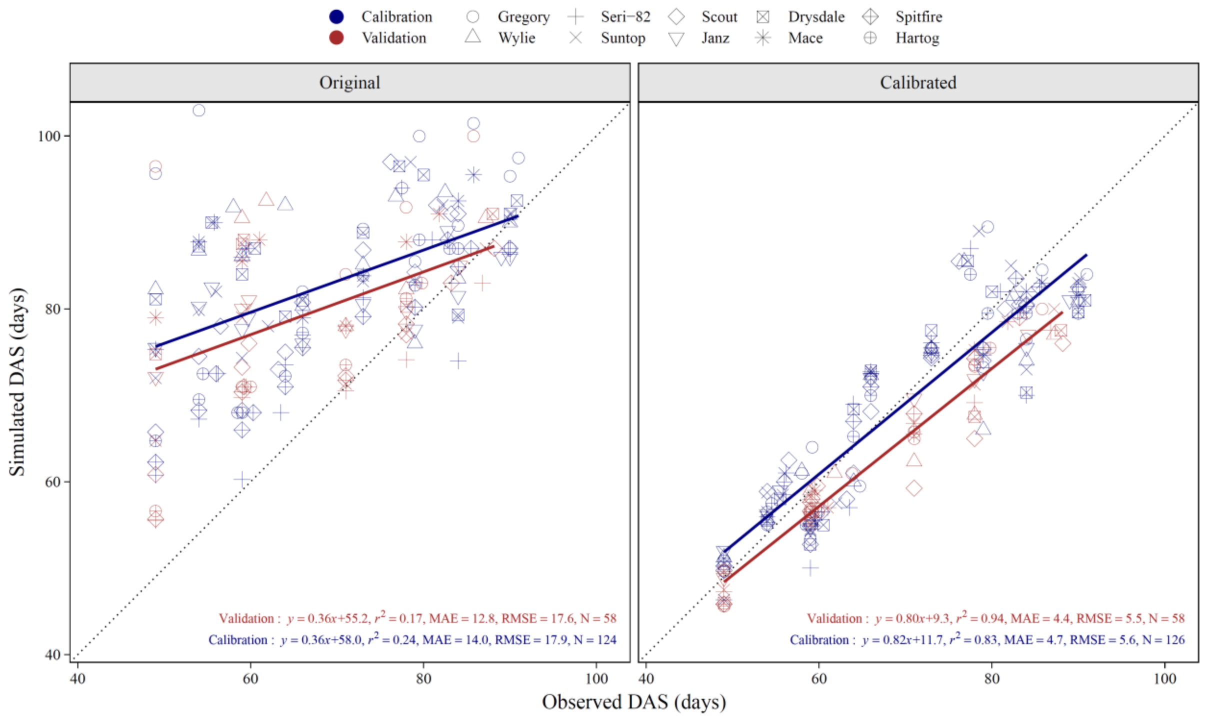


Figure S1. APSIM-wheat calibration results with the SUFI-2M algorithm across three locations in eastern Australia (south-eastern Queensland) with the parameters values from the APSIM default culvivar-specific parameter sets **(left)** and with re-calibrated parameters **(right)**. Data were averaged across replications in panel. RMSE is the root mean square error, MAE is the mean absolute error and N is the number of points. Observations included Zadoks growth stages from stem elongation (Z31) up to flowering (Z65).

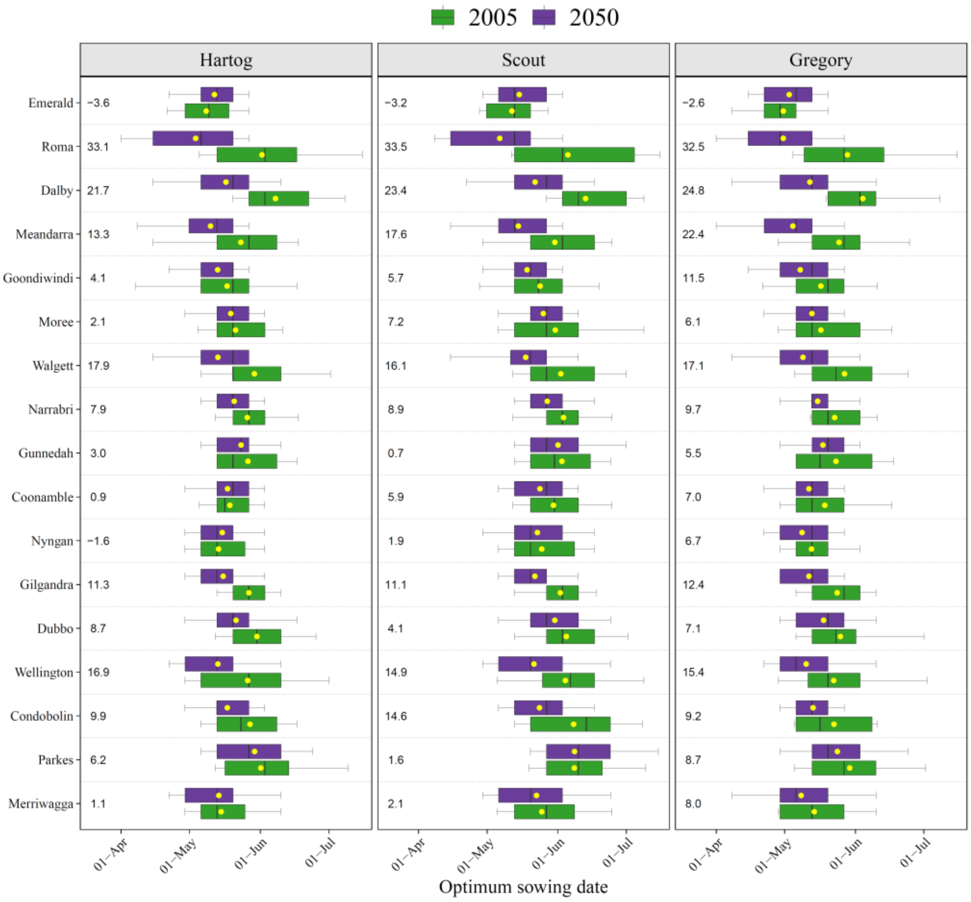


Figure S2. Sowing windows for the three selected spring wheat cultivars at the 17 sites in northeast Australia under the 2005 (current) and 2050 (future) climates. The numbers on the left of each panel show the changes in middle points of the sowing windows in 2050 relative to 2005. Positive values show earlier sowing dates in 2050 compared with 2005.

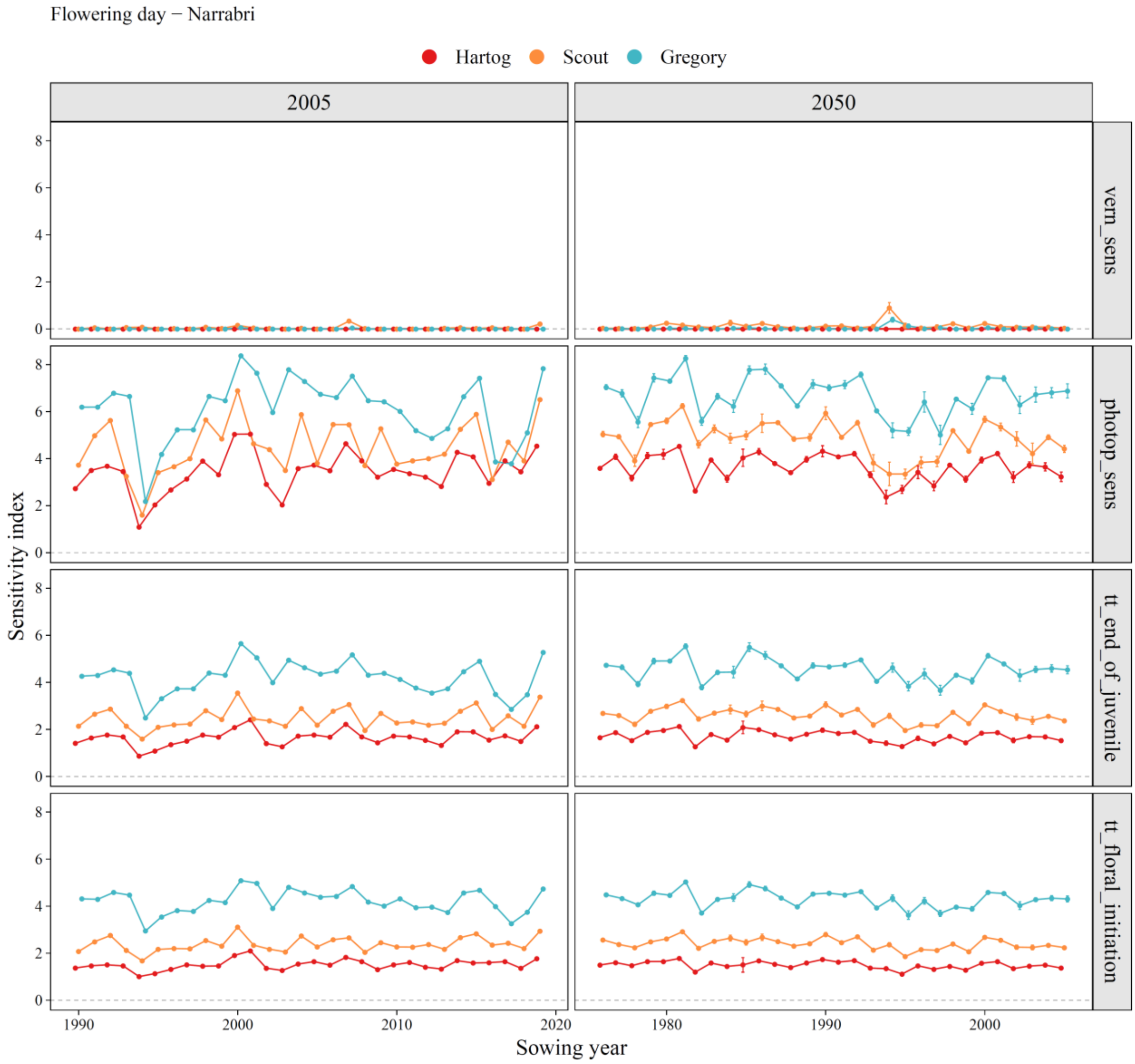


Figure S3. Sensitivity indices of the selected four phenological parameters for the simulated flowering day under the 2005 (current) and 2050 (future) climates for the three selected cultivars across the 30 simulated seaons in Narrabri. Parameters included: (1) vernalisation sensitivity (vern_sens), (2) photoperiod sensitivity (photop_sens), (3) thermal time from 'end of juvenile' to 'floral initiation' (tt_end_of_juvenile), and (4) thermal time from 'floral initiation' to 'flowering' (tt_floral_initiation).

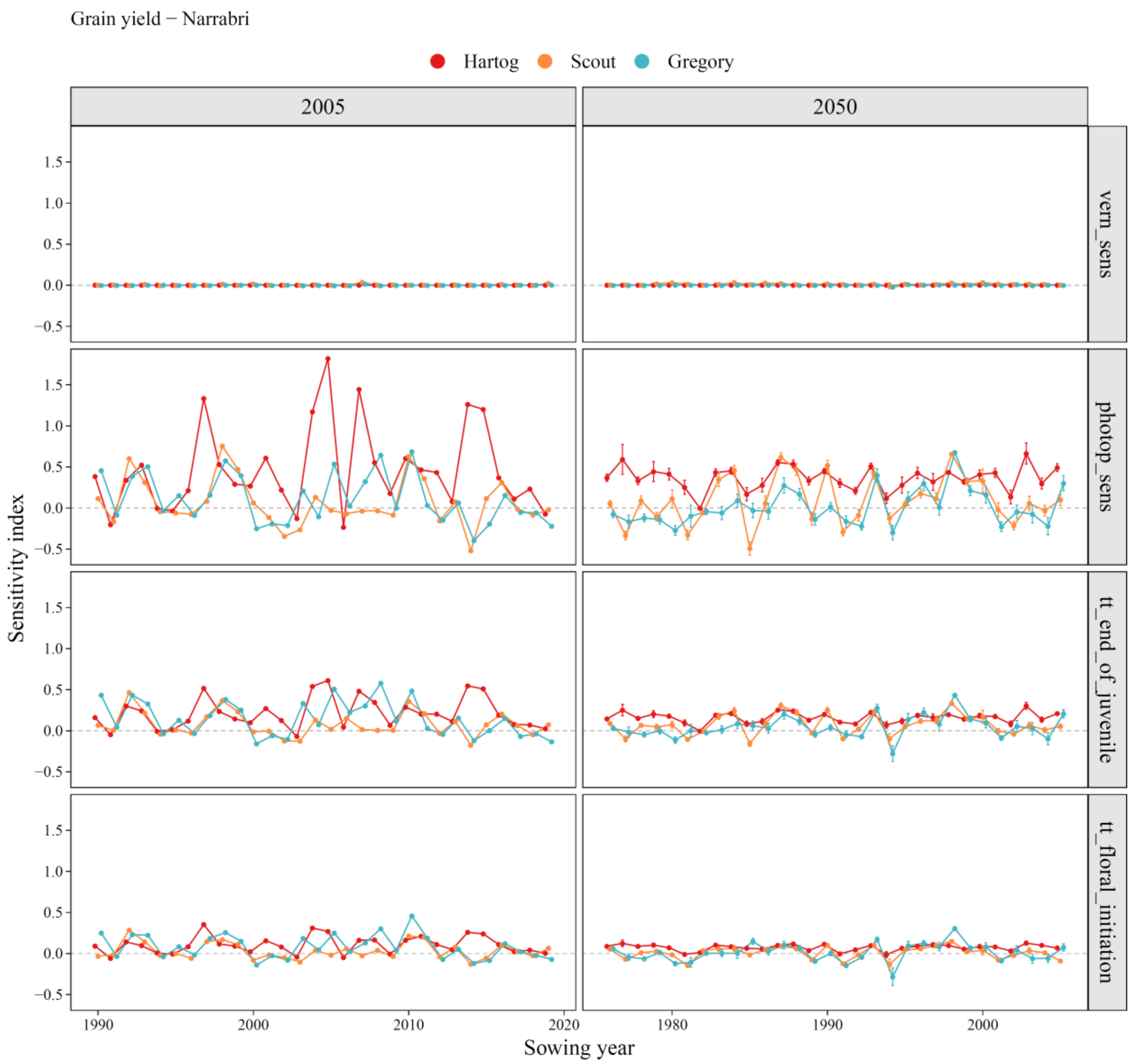


Figure S4. Sensitivity indices of the selected four phenological parameters for the simulated grain yield under the 2005 (current) and 2050 (future) climates for the three selected cultivars across the 30 simulated seaons in Narrabri. Parameters included: (1) vernalisation sensitivity (vern_sens), (2) photoperiod sensitivity (photop_sens), (3) thermal time from 'end of juvenile' to 'floral initiation' (tt_end_of_juvenile), and (4) thermal time from 'floral initiation' to 'flowering' (tt_floral_initiation).

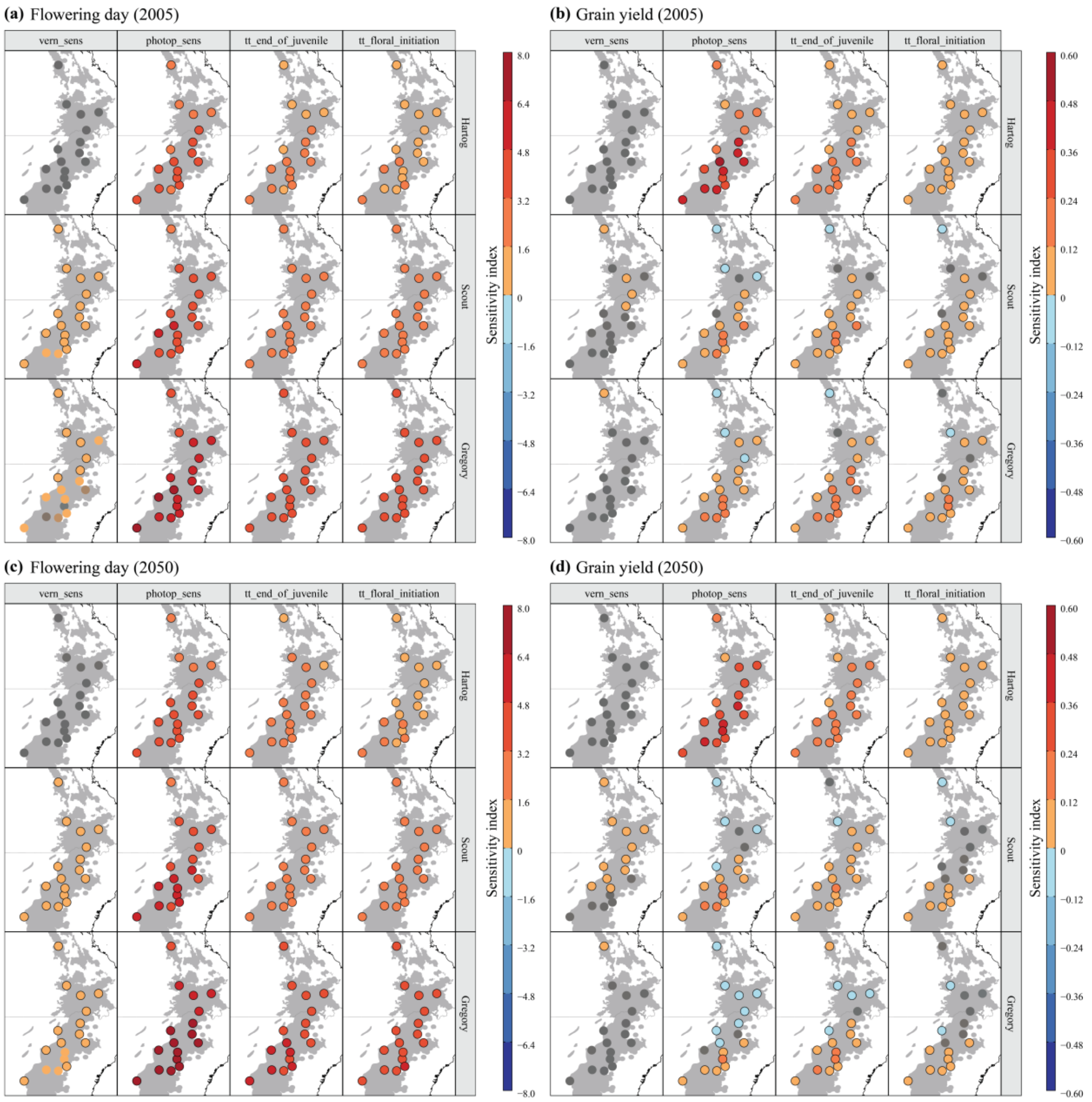


Figure S5. Sensitivity indices (averaged across 30 seasons) of the selected four phenological parameters related to flowering day **(a,c)** and grain yield **(b,d)** under the 2005 (current, **a-b**) and 2050 (future, **c-d**) climates for three selected spring wheat cultivars across 17 sites in eastern Australia. Parameters included: (1) vernalisation sensitivity (vern_sens), (2) photoperiod sensitivity (photop_sens), (3) thermal time from 'end of juvenile' to 'floral initiation' (tt_end_of_juvenile), and (4) thermal time from 'floral initiation' to 'flowering' (tt_floral_initiation).

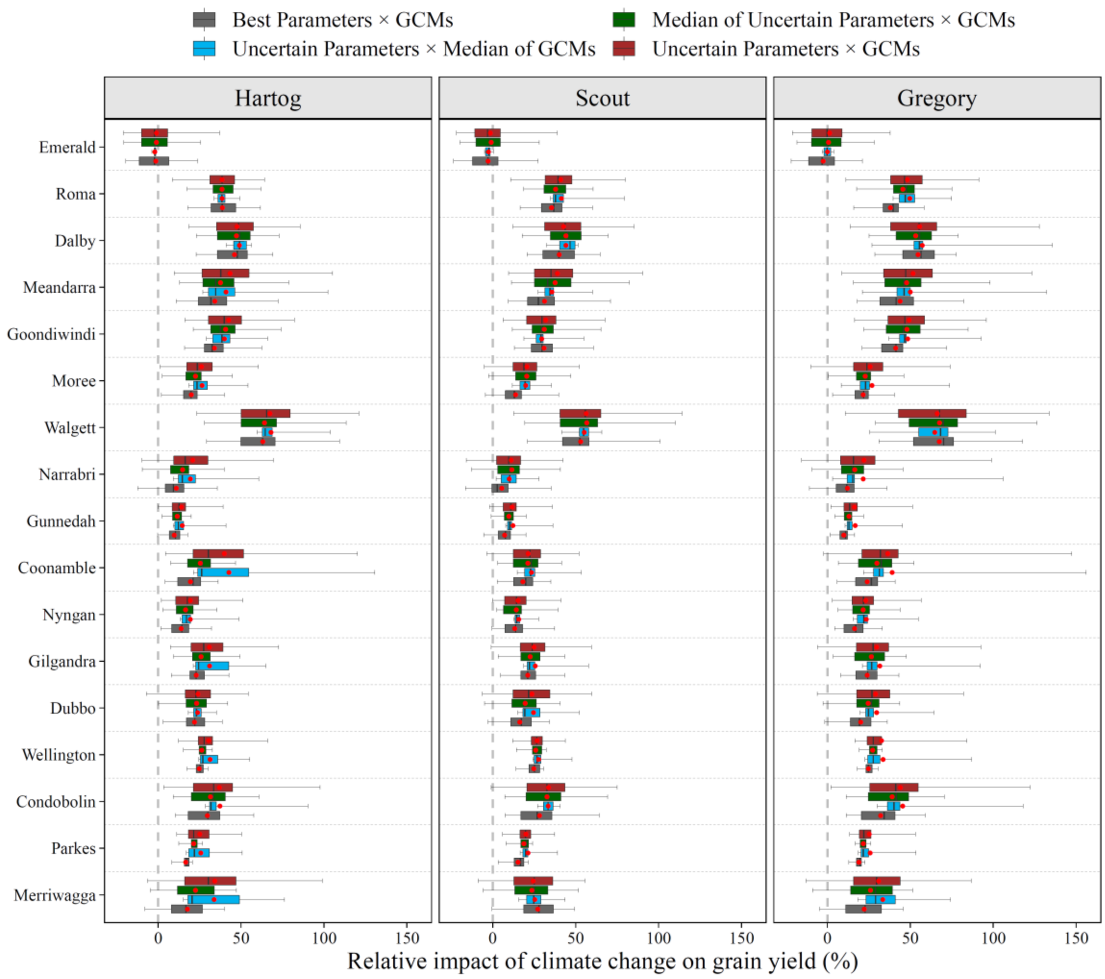


Figure S6. Impact of climate change on grain yield at the 17 selected sites in eastern Australia simulated with the best and uncertain parameter sets for three selected spring wheat cultivars. Box plots show the 10-25-50-75-90th quantiles along with the means (red points).
